# Genome Editing Is Induced in a Binary Manner in Single Human Cells

**DOI:** 10.1101/2022.03.04.482947

**Authors:** Gou Takahashi, Daiki Kondo, Minato Maeda, Yuji Morishita, Yuichiro Miyaoka

## Abstract

Even when precise nucleotide manipulations are intended, the outcomes of genome editing can be diverse, often including random insertions and deletions. The combinations and frequencies of these different outcomes in single cells are critical not only in the generation of genetically modified cell lines but also in the evaluation of the clinical effects of genome editing therapies. However, current methods only analyze cell populations, not single cells. Here, we utilized the Single Particle isolation System (SPiS) for the efficient isolation of single cells to systematically analyze genome editing results in individual human cultured cells. As a result, we discovered that genome editing induction has a binary nature, that is, the target alleles of cells tend to be all edited or not edited at all. This study enhances our understanding of the induction mechanism of genome editing and provides a new strategy to analyze genome editing outcomes in single cells.

## Introduction

Genome editing allows us to manipulate genetic information in basically any type of cell, and has been revolutionary in basic science, agriculture, and medicine^1–3^. Genome editing tools were originally designed to cleave target sequences in the genome DNA in the cell, so that genetic manipulations can be introduced into the genome via activated DNA repair pathways at the target sites^4^. These DNA repair pathways include non-homologous end-joining (NHEJ) and homology-directed repair (HDR)^5^. Each pathway gives distinct genome editing outcomes.

Although base editing and prime editing technologies do not require DNA double-strand breaks to manipulate the genome DNA sequence, they can still never produce only one type of editing^6–9^. Despite significant progress in the prediction of genome editing outcomes^10–13^, it has been impossible to produce a sole genetic manipulation. In other words, genome editing outcomes are always mixtures of different modifications of DNA sequences, such as insertions or deletions of different sizes and targeted recombination events. Therefore, for the application of genome editing, it is important to precisely measure its outcomes.

Various types of techniques have been used to analyze genome editing outcomes^14^, including sequencing-based methods (e.g., amplicon sequencing and TIDE^15^), denaturation-based methods (e.g., T7E1^16^ and single-stranded conformational polymorphism [SSCP] assays^17^), and digital PCR-based methods^18,19^. However, all of these methods are designed to analyze cell populations, not single cells. The combination of fluorescent reporter systems, such as traffic light reporter^20^ and flow cytometry, can visualize genome editing results in individual cells, but this setting cannot be applied to endogenous genes. To fully exploit the potential of genome editing, it is critical to grasp the editing outcomes in individual cells. For example, in a hypothetical situation where 50% of a population of diploid cells are WT/HDR heterozygotes and the other 50% are NHEJ/NHEJ homozygotes (Population 1 in Fig. 1a), and the other population consists of 50% of WT/NHEJ and 50% of HDR/NHEJ heterozygotes (Population 2 in Fig. 1a), the two populations would—as a whole—have identical allelic frequencies of WT, HDR, and NHEJ, while the cells would show a totally different composition (Fig. 1a). Therefore, there is a strong demand for an efficient strategy to investigate genome editing outcomes in single cells. The main reason why analyzing genome editing results in single cells has been so difficult is the limitation in the number of target molecules (i.e., a diploid cell has only two copies of genomic DNA). Even karyotypically abnormal cell lines only have several copies of target DNA sequences per cell at most. This is in clear contrast to the recent advancement of the single cell transcriptome^21^, where a typical mammalian cell has ~10^5^ mRNA molecules to analyze^22^.

**Figure 1.**
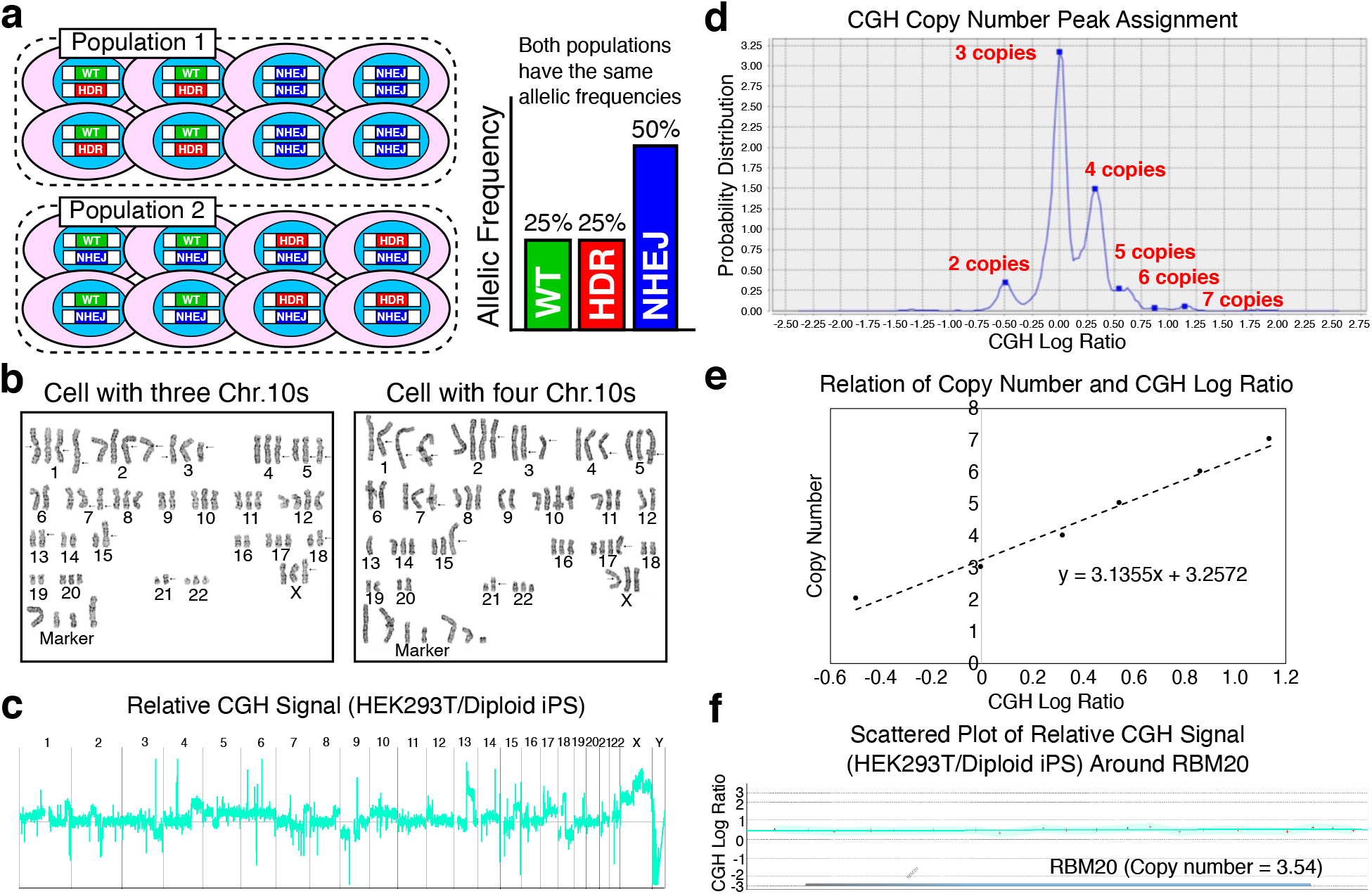
Measurement of the copy number of RBM20 in HEK293T cells. (a) A hypothetical situation that emphasizes the importance of the analysis of genome editing outcomes at the single cell level. Cell populations 1 and 2 consist of cells with totally different genotypes individually. However, the total allelic frequencies of WT, HDR, and NHEJ are exactly the same for both populations. (b) Representative karyotypes of HEK293T cells with three and four chromosome 10s. (c) A CGH analysis of HEK293T cells in comparison to diploid human iPS cells. The relative CGH signal of HEK293T cells normalized to that of diploid iPS cells is shown throughout the genome. (d) CGH copy number peak assignment. In the CGH analysis, there were several peaks of the CGH signal ratio between HEK293T cells and diploid iPS cells based on the chromosomal numbers. The highest peak corresponded to three copies per cell, and was set as the baseline of the CGH log ratio between HEK293T cells and iPS cells. (e) Line of fit between the copy number and the CGH log ratio based on the peak assignment shown in (d). The copy number and the CGH log ratio showed a clear linear correlation. (f) Scattered plot of relative CGH signal of HEK293T cells in comparison to diploid human iPS cells around the RBM20 gene. The relative CGH signals of HEK293T cells normalized by that of iPS cells are represented by +. No microduplications or microdeletions were detected around the RBM20 locus.

Therefore, it is extremely challenging to directly analyze the genomic DNA sequences in single cells. However, there are two clear advantages in the analysis of genomic DNA: one is that cells can replicate their own genome DNA as long as they are alive and proliferate; the other is that its sequence does not change, in contrast to the repertory of mRNA that changes dynamically depending on the cell state. Therefore, we decided to systematically isolate clones from a pool of genome edited cells. To accomplish this task, we utilized the Single Particle isolation System (SPiS) (On-chip Biotechnologies). The SPiS is an automated single cell dispenser that can gently and accurately dispense single cells in multi-well plates using image recognition technology to monitor the number of cells in aliquots^23,24^. The SPiS enabled us to isolate clones derived from genome edited cells with unprecedented efficiency. Our new pipeline to analyze genome editing based on the SPiS and findings from it will greatly contribute to improving the understanding of how genome editing occurs in the cell.

## Results

### Measurement of the copy number of RBM20 in HEK293T cells

First, we measured the copy number of our target gene, RBM20, per HEK293T cell to comprehend genome editing outcomes in single cells, because cell lines often have abnormal karyotypes. Therefore, we combined karyotyping and a comparative genomic hybridization (CGH) analysis to precisely estimate the copy number. We first karyotyped a total of 10 HEK293T cells, and found extensive chromosomal abnormalities (Fig. 1b). The most frequent chromosomal number was three (Supplementary Table 1). We also found out that five cells each had three and four chromosome 10s where RBM20 is located, respectively (Fig. 1b and Supplementary Fig. 1a).

Next, we analyzed the genome DNA of HEK293T cells by a CGH analysis. We compared HEK293T cells with WTC11 iPS cells that had been confirmed to be diploid^19^. The CGH analysis also demonstrated the chromosomal abnormalities of HEK293T cells, as we revealed by karyotyping (Fig. 1b, c). We detected peaks of the CGH signal corresponding to the chromosome number in HEK293T cells (Fig. 1d). Because the most frequent chromosome number was three (as determined by karyotyping), the most frequently observed peak of the CGH signal corresponded to three copies. We were able to incrementally assign copy numbers to these peaks to draw a line of fit between the relative CGH signal intensity and the copy number (Fig. 1d, e). By applying the relative median signal intensity of HEK293T cells to WTC11 iPS cells around the RBM20 gene (Fig. 1f and Supplementary Fig. 1b, c) to this line of fit, we estimated the average copy number of RBM20 to be 3.54, which matched the karyotyping results (Fig. 1b).

### Efficient isolation of HEK293T cell clones driven by the SPiS

To systematically analyze the genome editing outcomes in individual cells, we established an efficient pipeline to isolate cell clones that had gone through the genome editing process. Using CRISPR-Cas9 and a single strand DNA donor in HEK293T cells, as we described previously^18^, we introduced the RBM20 R636S mutation (c.1906C>A, chr.10), which causes inherited dilated cardiomyopathy^25^ (Fig. 2a). We used pX458 (Addgene plasmid #48138) to express EGFP via the T2A peptide fused to Cas9 in combination with gRNA. Therefore, the expression of EGFP guaranteed the expression of the CRISPR components in the same cell (Fig. 2b and Supplementary Fig. 2a). To minimize damage in the sorting of these EGFP-positive cells, we used a microfluidic cell sorter, On-chip Sort^26^. We confirmed that the sorting of EGFP-positive cells successfully enriched genome-edited cells (Supplementary Fig. 2b). These sorted cells were then subjected to the single cell isolation process using the SPiS, in which 384 cells were individually plated into four 96-well plates. The SPiS aspirates diluted cell suspension into a microtip and analyzes the images (taken by a CCD camera) of the contents. The camera takes two images with a one-second interval; thus, the contents in the microtip go down by gravity.

**Figure 2.**
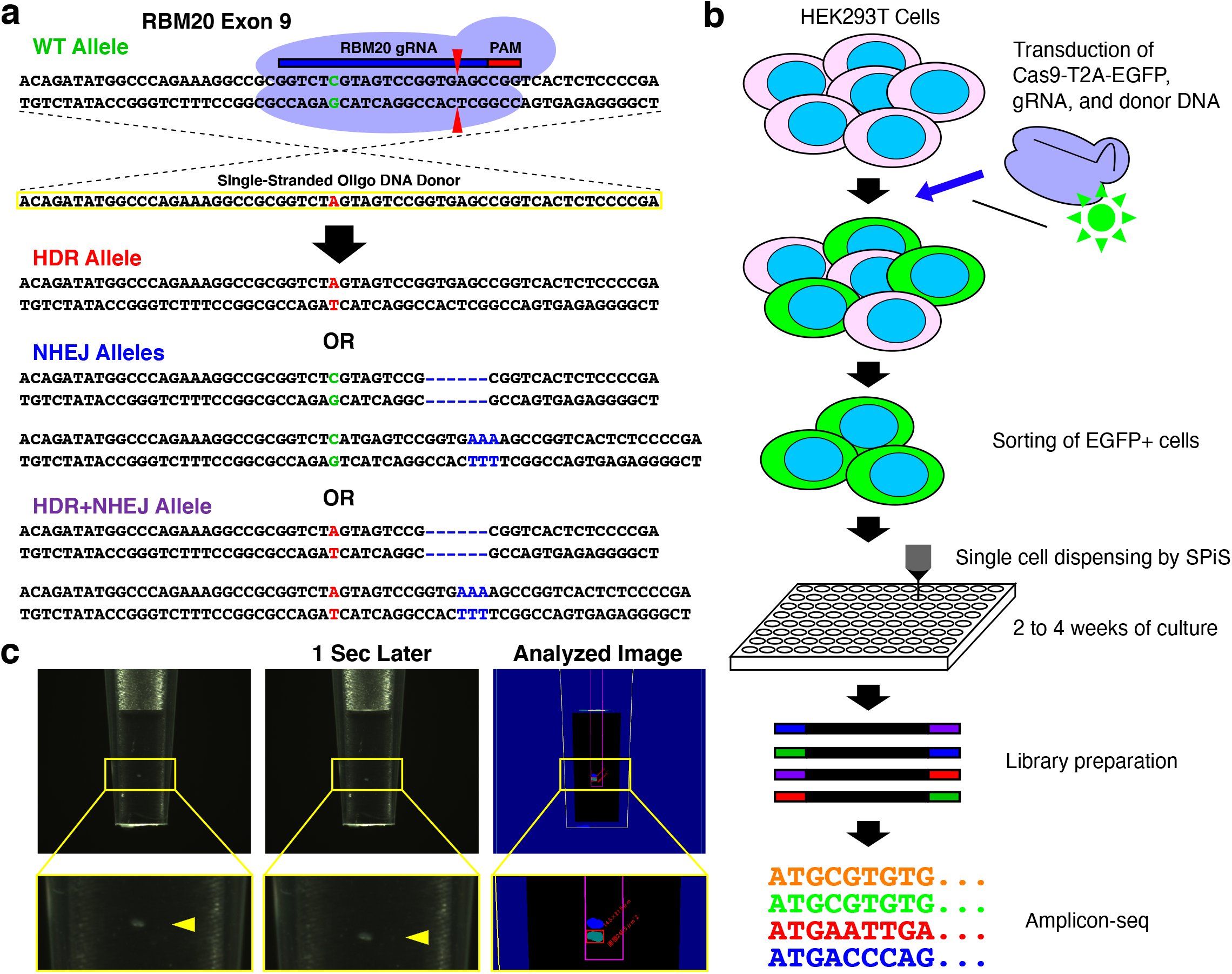
Schematic illustration of the experiments of this study. (a) Design of genome editing in RBM20 as a model case in this study. We introduced the R636S (c.1906C>A) mutation using CRISPR-Cas9 and a single-stranded oligonucleotide donor DNA. The resulting HDR allele has a C to A single nucleotide substitution, whereas the NHEJ alleles have various insertions and deletions. (b) The experimental flow of this study. We expressed Cas9-T2A-EGFP together with the gRNA targeting RBM20 to label HEK293T cells in which Cas9 protein was expressed. Using a microfluidic cell sorter (On-chip Sort), we sorted EGFP+ cells. Then, the sorted cells were plated into four 96-well plates by the SPiS. Cells were cultured for 2 to 4 weeks and then the genome editing outcomes were analyzed by amplicon sequencing. (c) An imaging analysis of single cells dispensed by the SPiS. The SPiS aspirates cell suspension into a specialized microtip and takes two images with a one-second interval. Only when the SPiS recognized a single cell, the content of the tip was dispensed into a well of a 96-well plate. Otherwise, the SPiS would take another aliquot of the cell suspension to repeat the process.

The SPiS analyzes the two images to measure the velocity of precipitation and the size of the contents (Fig. 2c). By doing this, the SPiS can discriminate cells from other objects (e.g., cell debris and dust). The SPiS dispenses contents of a microtip into a culture dish, only when the system recognizes one cell. After two to four weeks of culture, genomic DNA of HEK293T cell clones derived from single cells was harvested for amplicon sequencing to analyze the outcomes of genome editing (Fig. 2b).

### Successful analysis of genome editing outcomes in single HEK293T cells

We conducted single cell isolation processes using the SPiS three times (Cas9 No.1 to No. 3 experiments), and obtained >90 clones in each attempt (Fig. 3a). We performed amplicon sequencing on the RBM20 target site and analyzed the outcomes in isolated clones using CRISPResso2^27^ (Supplementary Fig. 3a). We defined the clean RBM20 R636S (c.1906C>A) substitution and insertions or deletions as “HDR” and “NHEJ” events, respectively, while combinations of both in the same allele were defined as “HDR+NHEJ” events (Fig. 2a and Supplementary Fig. 3b). HDR+NHEJ events occurred at a relatively low frequency in comparison to NHEJ, and the HDR frequency was even lower (Fig. 3a). Based on these HEK293T cell clones with edited alleles, we were able to estimate the overall allelic frequencies of each genome editing event (i.e., WT, NHEJ, HDR, and HDR+NHEJ) (Fig. 3b). Thus, our SPiS-based approach enables us to systematically analyze genome editing outcomes in individual cells.

**Figure 3.**
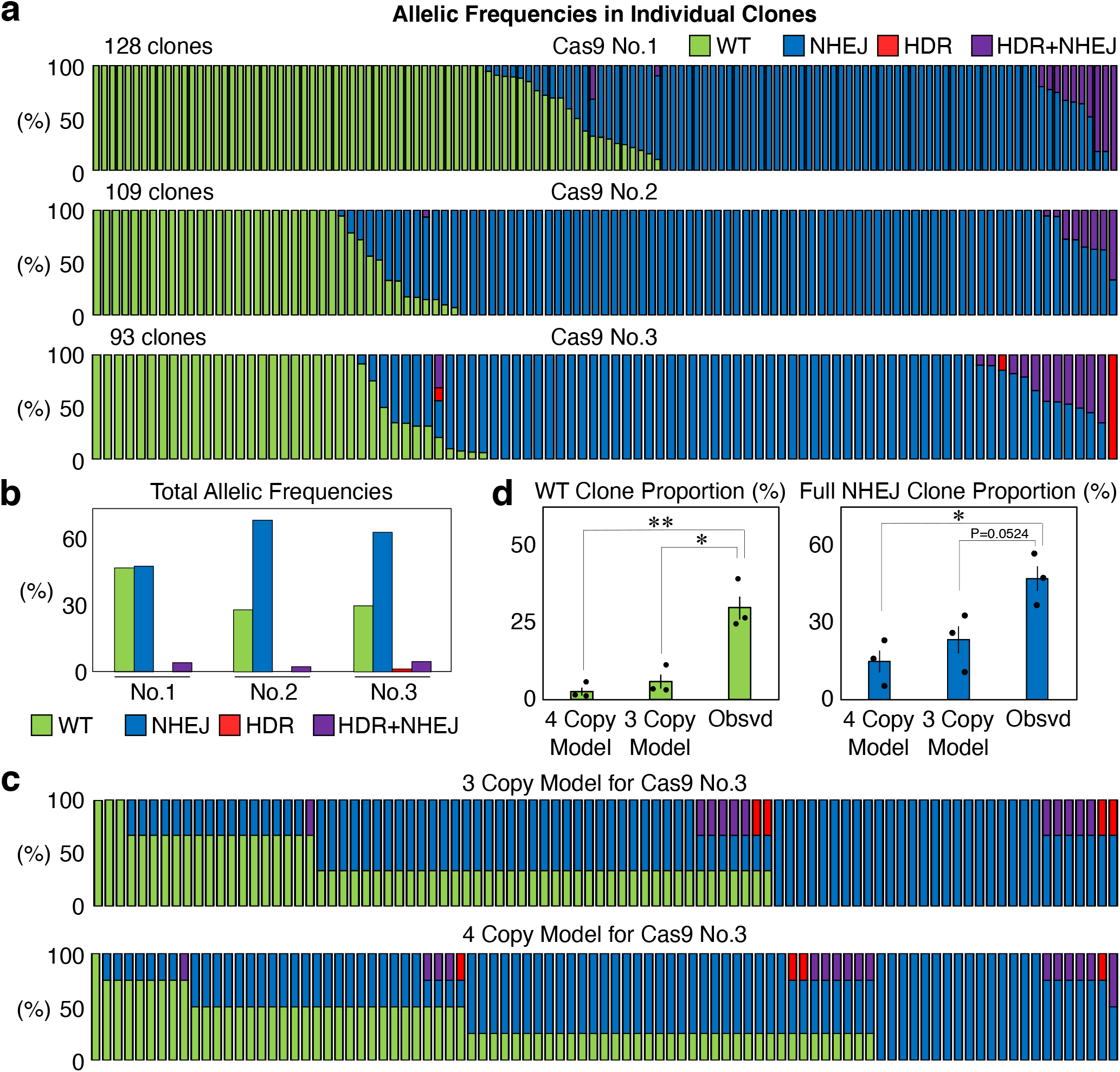
Genome editing outcomes in RBM20 in individual HEK293T cells edited by Cas9. (a) Genome editing outcomes in isolated clones derived from single HEK293T cells edited by Cas9. We repeated the experiment three times, and isolated more than 90 clones out of 384 cells plated in all three trials. Each bar represents one clone, and the proportions of WT (green), NHEJ (blue), HDR (red), and HDR+NHEJ (purple) in one clone are also shown in each bar. (b) Total allelic frequencies in genome edited HEK293T cells in the three experiments shown in (a). (c) Models of the distributions of HEK293T cell clones with different genome editing outcomes in the Cas9 No.3 experiment, if genome editing randomly occurred at the frequencies shown in (b) in HEK293T cells with three or four copies of RBM20. (d) Comparison of the proportions of WT and full NHEJ clones between the mathematical models with three and four copies of RBM20 and the actually observed cells. Values ± S.E. are shown (n=3). Student’s t-test was used to evaluate differences. *P<0.05 and **P<0.01.

### Binary induction of genome editing in single cells

One notable feature in our results was that roughly half of the cells had all their target RBM20 alleles edited by NHEJ (47/128, 62/109, and 44/93, respectively). On the other hand, there were quite a few cells that remained completely unedited (49/128, 26/109, and 24/93, respectively), although these cells had expressed Cas9 and gRNA (Fig. 3a). These observations were quite surprising considering the fact that these cells have three or four copies of RBM20 on average (Fig. 1).

To address whether this trend is significant, we mathematically calculated the number of HEK293T cell clones with different genome editing outcomes assuming that these events occurred randomly. Because the copy number of RBM20 was 3.54 (Fig. 1), we built two models with three or four copies of RBM20 (Fig. 3c and Supplementary Fig. 4). In these models, we calculated how many cells with three or four copies of RBM20 should have specific genotypes based on the overall frequencies of WT, NHEJ, HDR, and HDR+NHEJ alleles. We compared the proportions of clones that are expected to be unedited to remain as WT or fully edited by NHEJ between the two models and the actual observation (Fig. 3a, c). As we expected, the proportions of actually isolated clones were significantly higher than those based on the models for both WT and full NHEJ clones (Fig. 3d). In addition, among 330 clones, we observed one clone in which all target alleles were modified by HDR (Fig. 3a). No such clone was expected in the mathematical models (Fig. 3c and Supplementary Fig. 4). These results suggested that genome editing occurs in a binary manner, in which cells show either no editing at all or full editing.

### HypaCas9 induces HDR more efficiently than Cas9 in single cells

Because the frequency of HDR was so low with Cas9, we were not able to investigate how HDR is induced in single cells (Fig. 3). HDR is often more desirable than NHEJ, as HDR can introduce precise manipulation of the DNA sequence. Therefore, we decided to apply our SPiS- based analysis to genome editing conditions that are more favorable for the induction of HDR. We found that Cas9 with improved proof-reading (e.g., HypaCas9^28^, Cas9-HF1^29^, and eSpCas9^30^) induced more HDR than Cas9 in our previous cell population-based assay^31^; thus, we targeted the same RBM20 R636S mutation in HEK293T cells using HypaCas9, and conducted the SPiS-based single cells assay three times. We confirmed that HypaCas9 produced more clones with HDR than Cas9 (Fig. 4a). We calculated the allelic frequencies of WT, NHEJ, HDR, and HDR+NHEJ (Fig. 4b), and compared these between Cas9 and HypaCas9. We found that the HDR allelic frequency was higher with HypaCas9 than with Cas9; however, the WT, NHEJ, and HDR+NHEJ allelic frequencies were comparable (Fig. 4c). Based on these overall allelic frequencies, we built mathematical models of HEK293T cell clones with genome editing by HypaCas9 in the same way as for Cas9 (Fig. 4d and Supplementary Fig. 5). The binary manner of genome editing induction was also observed with HypaCas9, as the WT and full NHEJ clone proportions were significantly higher than the mathematical models (Fig. 4e). Moreover, binary induction was also observed in HDR. Among 422 clones, we isolated a total of six clones with all alleles edited by HDR (Fig. 4a). Because the overall frequency of HDR was 5.89% (Fig. 4c), no such clone was expected to be isolated if genome editing events occurred randomly in single cells (1 out of 4894 cells would be expected to have having three HDR alleles and 1 out of 83088 cells would be expected to have four HDR alleles). Thus, we confirmed the binary induction of genome editing, even with HypaCas9, which induces more HDR than Cas9. The binary mode was observed in the induction of both HDR and NHEJ.

**Figure 4.**
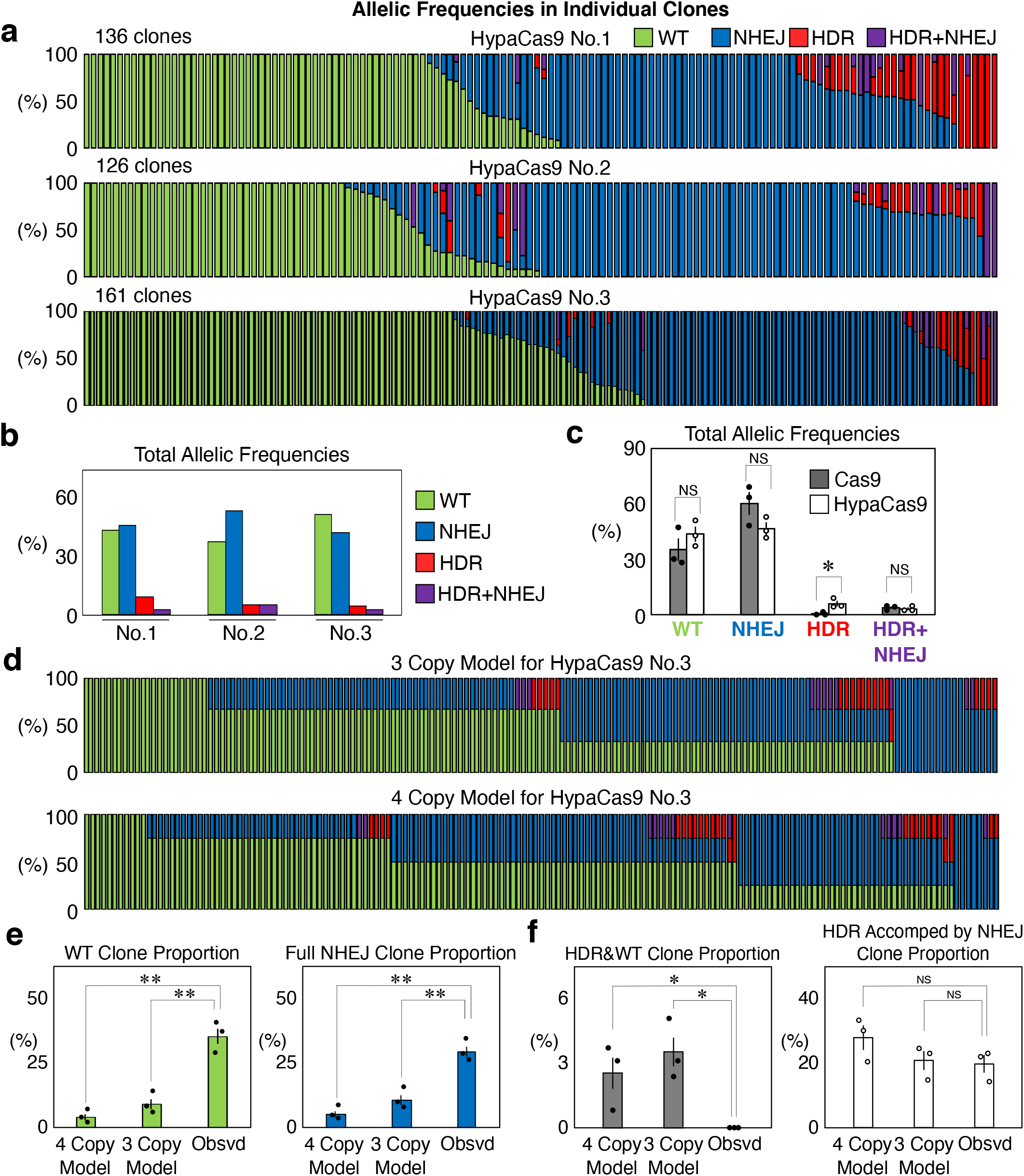
Genome editing outcomes in RBM20 in individual HEK293T cells edited by HypaCas9. (a) Genome editing outcomes in isolated clones derived from single HEK293T cells edited by HypaCas9 and the single-stranded donor DNA. We repeated the same experiment three times. Each bar represents one clone, and the proportions of WT (green), NHEJ (blue), HDR (red), and HDR+NHEJ (purple) in one clone are also shown in each bar. (b) Total frequencies of WT (green), NHEJ (blue), HDR (red), and HDR+NHEJ (purple) allelic frequencies in genome edited HEK293T cells in the three experiments shown in (a). (c) Comparison of total allelic frequencies of WT, NHEJ, HDR, and HDR+NHEJ between Cas9 and HypaCas9. Values ± S.E. are shown (n=3). Student’s *t*-test was used to evaluate differences. *P<0.05 and NS: not significantly different (P>0.1). (d) Models of the distributions of HEK293T cell clones with different genome editing outcomes in the HypaCas9 No.3 experiment, if genome editing randomly occurred at the frequencies shown in (b) in HEK293T cells with three or four copies of RBM20. (e) Comparison of the proportions of WT and full NHEJ clones between the mathematical models with three and four copies of RBM20 and the actually observed cells. Values ± S.E. are shown (n=3). Student’s *t*-test was used to evaluate differences. **P<0.01. (f) Comparison of the proportions of clones with partial editing by HDR and clones with HDR accompanied by NHEJ events between the models with three and four copies of RBM20 and the actually observed cells. Values ± S.E. are shown (n=3). Student’s *t*-test was used to evaluate differences. *P<0.05 and NS: not significantly different (P>0.1).

### HDR is more often accompanied by NHEJ than WT

We noticed that clones with HDR often had NHEJ alleles unless cells were fully edited by HDR. Therefore, we calculated the proportions of clone that would be expected to have only HDR and WT alleles, as these clones represent partial editing by HDR. Even though 2.5–3.5% of the cells were expected to have only HDR and WT alleles, we isolated no such clones (Fig. 4a, f). We further investigated the expected and observed the proportions of clones with both HDR and NHEJ events, including any clones with the HDR+NHEJ alleles, as they represent HDR and NHEJ events induced in the same alleles. As we expected, the proportions of clones with both HDR and NHEJ events that were observed and those that were predicted by the models were comparable (Fig. 4f). These results coincide with the fact that cells tend to have all target alleles edited when genome editing was induced, so partial editing is rare (Fig. 4e). Moreover, HDR and NHEJ are often induced together in single cells (Fig. 4f).

### Genome editing is also induced in a binary manner in ATP7B and GRN

We next addressed whether the observed binary fashion of genome editing stands in other target sites. Therefore, we targeted ATP7B and GRN to introduce R778L (c.2333G>T, chr.13)^32^ and R493X (c.1477C>T, chr.17)^33^ mutations, respectively into HEK293T cells as described previously^18^. HypaCas9 and single stranded donor DNA were used in the same way as for RBM20 R636S mutagenesis (Supplementary Fig. 6a, b). We isolated clones and performed amplicon sequencing to quantify the frequency of genome editing (Fig. 5a-e). To build models for genome editing of ATP7B and GRN, we quantified the copy numbers of these genes, which were determined to be 2.88 for ATP7B and 3.54 for GRN (Fig. 1 and Supplementary Fig. 6c, d). We then calculated the expected clone numbers with the assumption that genome editing events occurred randomly in the cells to build models for genome editing of ATP7B and GRN (Supplementary Fig. 7a, b), and compared them to the observed results. Similar to the RBM20 R636S mutagenesis, the number of clones in which all target alleles were edited by NHEJ or not edited at all was significantly higher in comparison to the models (Fig. 5b, e). We also isolated clones in which all target alleles were edited by HDR (8/280 and 2/276 for ATP7B and GRN, respectively) (Fig. 5a, d). This was particularly noteworthy for GRN, as the overall efficiency of HDR induction was only 1.02% (Fig. 5d). Collectively, these results demonstrate that the binary nature of genome editing is not restricted to a specific target site. Moreover, we did not isolate any clones with partial HDR editing in ATP7B or GRN; however, HDR and NHEJ were often induced together in single cells, similarly to in RBM20 (Figs. 5a, c, d).

**Figure 5.**
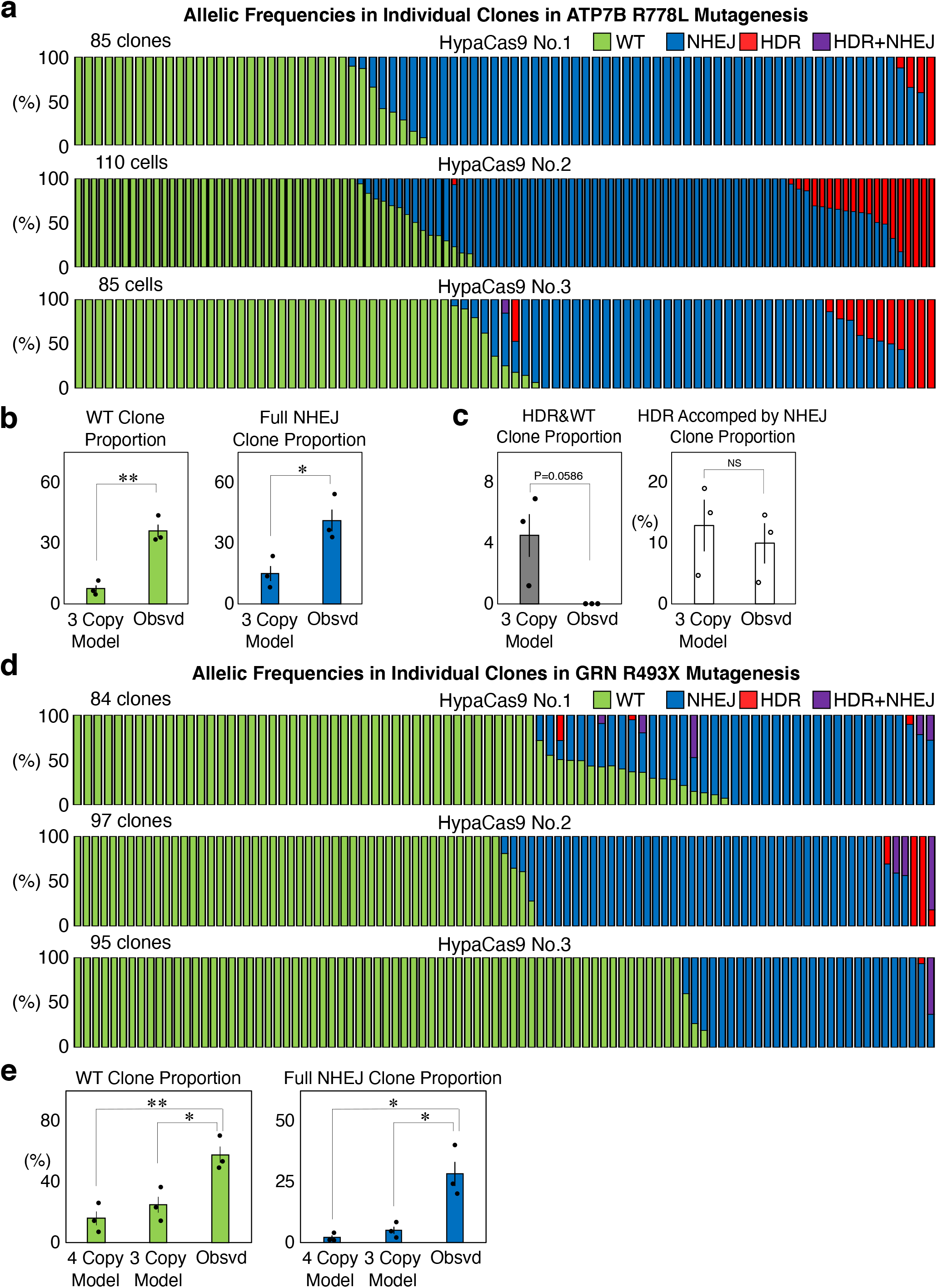
Genome editing outcomes in ATP7B and GRN in individual HEK293T cells edited by HypaCas9. (a) Genome editing outcomes in isolated clones derived from single HEK293T cells edited by HypaCas9 and the single-stranded donor DNA targeting ATP7B. We repeated the same experiment three times. Each bar represents one clone, and the proportions of WT (green), NHEJ (blue), HDR (red), and HDR+NHEJ (purple) in one clone are also shown in each bar. (b) Comparison of the proportions of WT and full NHEJ clones between the mathematical models with three copies of ATP7B and the actually observed cells. Values ± S.E. are shown (n=3). Student’s *t*-test was used to evaluate differences. *P<0.05 and **P<0.01. (c) Comparison of the proportions of clones with partial editing by HDR and clones with HDR accompanied by NHEJ events between the models with three copies of ATP7B and the actually observed cells. Values ± S.E. are shown (n=3). Student’s *t*-test was used to evaluate differences. NS: not significantly different (P>0.1). (d) Genome editing outcomes in isolated clones derived from single HEK293T cells edited by HypaCas9 and the single-stranded donor DNA targeting GRN. We repeated the same experiment three times. Each bar represents one clone, and proportions of WT (green), NHEJ (blue), HDR (red), and HDR+NHEJ (purple) in one clone are also shown in each bar. (e) Comparison of the proportions of WT and full NHEJ clones between the mathematical models with three and four copies of GRN and the actually observed cells. Values ± S.E. are shown (n=3). Student’s *t*-test was used to evaluate differences. *P<0.05 and **P<0.01.

### Binary genome editing is also prominent in HeLa cells

We further addressed whether the binary induction of genome editing occurs in other cell types. For this purpose, we introduced the RBM20 R636S (c.1906C>A), ATP7B R778L (c.2333G>T), and GRN R493X (c.1477C>T) mutations into HeLa cells in the same way as for HEK293T cells. We were able to apply our SPiS-based system to isolated HeLa cell clones that had gone through the genome editing process, although HeLa cells exhibited lower cell survival in comparison to HEK293T cells (Fig. 6a-g). We measured the copy numbers of these target genes by combining the karyotyping and CGH analyses in HeLa cells (Supplementary Fig. 8 and Supplementary Table 2). Based on these copy numbers, mathematical models of clones with different genotypes were built in the same way as for HEK293T cells (Supplementary Fig. 9).

**Figure 6.**
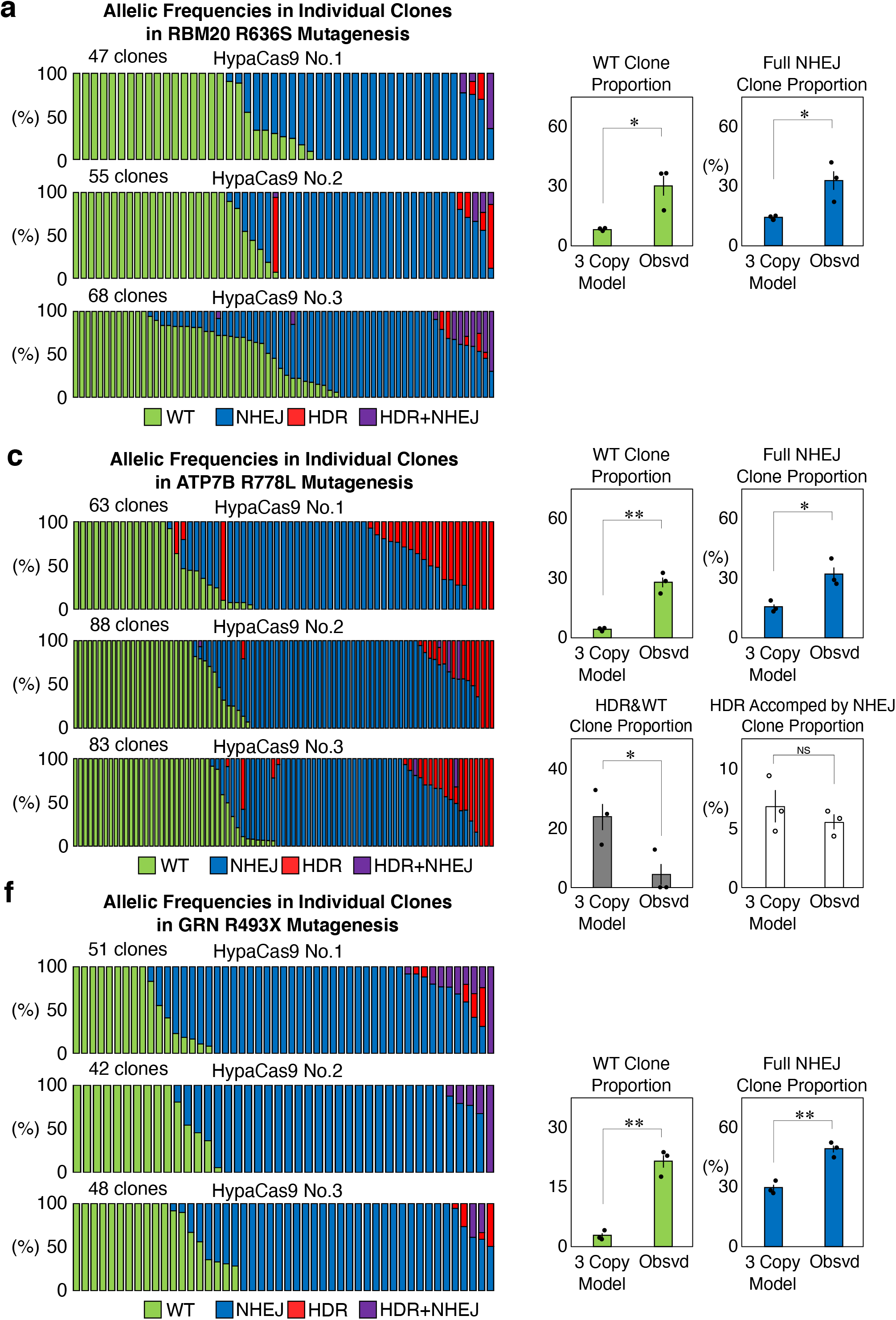
Genome editing outcomes in individual HeLa cells. (a) Genome editing outcomes in isolated clones derived from single HeLa cells edited by HypaCas9 and the single-stranded donor DNA targeting RBM20. Each bar represents one clone, and the proportions of WT (green), NHEJ (blue), HDR (red), and HDR+NHEJ (purple) in one clone are also shown in each bar. (b) Comparison of the proportions of WT and full NHEJ clones between the mathematical models with three copies of RBM20 and the actually observed cells. Values ± S.E. are shown (n=3). Student’s *t*-test was used to evaluate differences. *P<0.05. (c) Genome editing outcomes in isolated clones derived from single HeLa cells edited by HypaCas9 and the single-stranded donor DNA targeting ATP7B. Each bar represents one clone, and the proportions of WT (green), NHEJ (blue), HDR (red), and HDR+NHEJ (purple) in one clone are also shown in each bar. (d) Comparison of the proportions of WT and full NHEJ clones between the mathematical models with three copies of ATP7B and the actually observed cells. Values ± S.E. are shown (n=3). Student’s *t*-test was used to evaluate differences. *P<0.05 and **P<0.01. (e) Comparison of the proportions of clones with partial editing by HDR and clones with HDR accompanied by NHEJ events between the models with three copies of ATP7B and the actually observed cells. Values ± S.E. are shown (n=3). Student’s *t*-test was used to evaluate differences. *P<0.05. NS: not significantly different (P>0.1). (f) Genome editing outcomes in isolated clones derived from single HeLa cells edited by HypaCas9 and the single-stranded donor DNA targeting GRN. Each bar represents one clone, and the proportions of WT (green), NHEJ (blue), HDR (red), and HDR+NHEJ (purple) in one clone are also shown in each bar. (g) Comparison of the proportions of WT and full NHEJ clones between the mathematical models with three copies of GRN and the actually observed cells. Values ± S.E. are shown (n=3). Student’s t-test was used to evaluate differences. **P<0.01.

We found that the observed proportions of clones that were unedited or fully edited by NHEJ were significantly higher than in the calculated models based on the overall allelic frequencies in all three genes (Fig. 6b, d, g). The HDR frequency was higher in ATP7B than in RBM20 or GRN in HeLa cells, and we isolated 10 clones fully edited by HDR out of 234 clones (Fig. 6c). In ATP7B, the proportion of clones with partial editing by HDR was significantly lower in comparison to the model, while that of clones with HDR accompanied by NHEJ was comparable to the model (Fig. 6e). These results indicate that the binary nature of genome editing induction is shared between HeLa cells and HEK293T cells.

### Binary genome editing is less evident in PC9 cells

Finally, we investigated the binary nature of genome editing in another cell line, PC9 cells. We introduced the same three mutations into PC9 cells and conducted our SPiS-based analysis (Fig. 7a-f). We noticed that the genome editing efficiency in PC9 cells was generally lower than that in HEK293T cells or HeLa cells, and that HDR was barely induced (Fig. 7a, c, e). We measured the copy numbers of the target genes in PC9 cells (Supplementary Fig. 10 and Supplementary Table 3). Based on these copy numbers, the mathematical models of clones with different genotypes were built for PC9 cells (Supplementary Fig. 11). We compared the proportion of clones that were unedited or fully edited by NHEJ between the observations and the models. We found that the trend regarding the binary induction of genome editing was noticeable but less evident, as the proportions of clones fully edited by NHEJ in RBM20 and clones unedited in ATP7B were the only significant differences between the observed clones and the models (Fig. 7b, d, f). Therefore, binary induction is a general feature of genome editing; however, the extent of binary induction can vary in different contexts.

**Figure 7.**
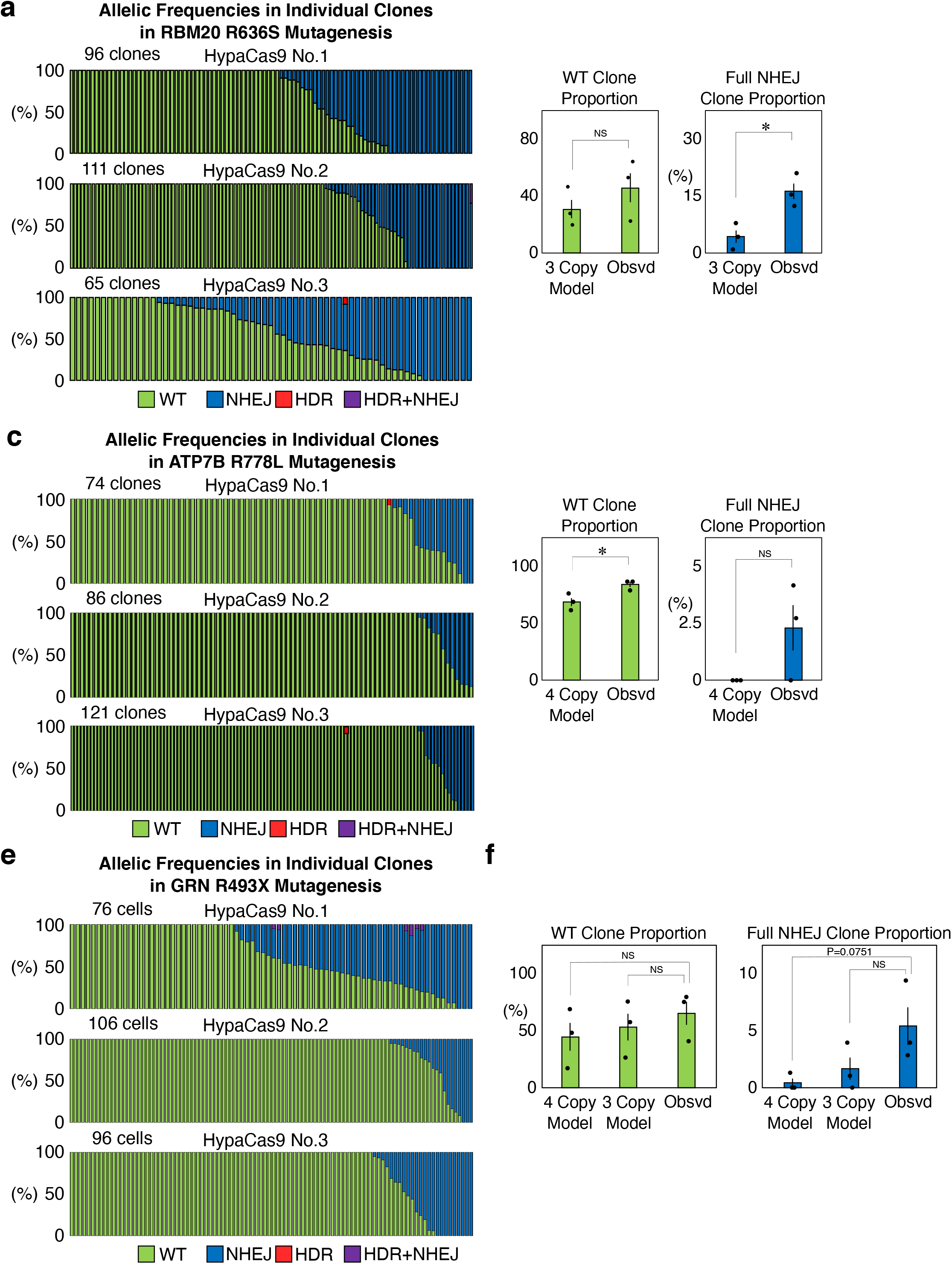
Genome editing outcomes in individual PC9 cells. (a) Genome editing outcomes in isolated clones derived from single PC9 cells edited by HypaCas9 and the single-stranded donor DNA targeting RBM20. Each bar represents one clone, and the proportions of WT (green), NHEJ (blue), HDR (red), and HDR+NHEJ (purple) in one clone are also shown in each bar. (b) Comparison of the proportions of WT and full NHEJ clones between the mathematical models with three copies of RBM20 and the actually observed cells. Values ± S.E. are shown (n=3). Student’s *t*-test was used to evaluate the difference. *P<0.05. NS: not significantly different (P>0.1). (c) Genome editing outcomes in isolated clones derived from single PC9 cells edited by HypaCas9 and the single-stranded donor DNA targeting ATP7B. Each bar represents one clone, and the proportions of WT (green), NHEJ (blue), HDR (red), and HDR+NHEJ (purple) in one clone are also shown in each bar. (d) Comparison of the proportions of WT and full NHEJ clones between the mathematical models with three copies of ATP7B and the actually observed cells. Values ± S.E. are shown (n=3). Student’s *t*-test was used to evaluate differences. *P<0.05. NS: not significantly different (P>0.1). (e) Genome editing outcomes in isolated clones derived from single PC9 cells edited by HypaCas9 and the single-stranded donor DNA targeting GRN. Each bar represents one clone, and the proportions of WT (green), NHEJ (blue), HDR (red), and HDR+NHEJ (purple) in one clone are also shown in each bar. (f) Comparison of the proportions of WT and full NHEJ clones between the mathematical models with three copies of GRN and the actually observed cells. Values ± S.E. are shown (n=3). Student’s *t*-test was used to evaluate differences. NS: not significantly different (P>0.1).

## Discussion

Most of the current methods used to analyze genome editing outcomes deal with cell populations. In this study, we were able to isolate single cell clones with an unprecedented efficiency by the SPiS. The SPiS not only increased the number of clones that we were able to isolate, but also ensured that the isolated clones were derived from single cells based on the image analysis, which avoided plating multiple cells together. This original system allowed us to investigate genome editing outcomes in single cells.

We were surprised to observe that most HEK293T cells were either unedited WT or had all targeted alleles fully manipulated by Cas9. The number of partially edited cells that had both unedited WT alleles and edited alleles was limited (Fig. 3). We previously reported that Cas9 variants with enhanced proof-reading are more efficient at inducing HDR than regular Cas9^31^. We confirmed that HypaCas9 produced more HDR in single cells than Cas9, and found that HypaCas9 also induced genome editing in a binary fashion (Fig. 4d). This trend was shared among all RBM20, ATP7B, and GRN genes in both HEK293T cells and HeLa cells, and to a lesser extent in PC9 cells (Figs. 4–7). Because we sorted cells that expressed EGFP together with Cas9 and gRNA, these WT clones escaped from DNA cleavage by Cas9, or precisely repaired the genomic DNA after DNA cleavage. We speculate that each cell has an intracellular environment that is favorable for a particular genome editing outcome, so that all target alleles in one cell tend to have the same fate. The average copy numbers of RBM20, ATP7B, and GRN in HEK293T cells, HeLa cells, and PC9 cells were 2.88-3.71 (Fig. 1 and Supplementary Figs. 1, 8, and 10). The fact that some cells have full HDR or HDR+NHEJ events in roughly 3 to 4 copies despite the much lower overall frequencies also suggests that target alleles in one cell tend to have the same fate (Figs. 4–6). This binary nature of genome editing can be beneficial if homozygous mutagenesis is necessary. Indeed, we previously reported the isolation of a human iPS cell line with a homozygous PHOX2B Y14X mutation generated by TALENs, even though the overall induction efficiency of HDR was just approximately 1%^19^. At the same time, however, it makes heterozygous mutagenesis extremely difficult, as we observed that HDR was more often accompanied by NHEJ rather than unedited WT alleles (Figs. 4–6).

At this point, however, the factors that determine which DNA repair pathway a single cell takes remain unknown. One possibility is that the cells respond differently to Cas9 cleavage at different points in the cell cycle. It is known that the frequency of HDR and NHEJ increases in the S/G2 and G1 phases, respectively^34,35^. Therefore, it would be interesting to address whether the cell cycle also influences this binary choice between full editing and no editing in single cells. PC9 cells showed a much lower efficiency of overall induction of genome editing than HEK293T cells and HeLa cells (Fig. 7). The binary fashion of induction of genome editing was also less evident in PC9 cells than in HEK293T cells and HeLa cells. This could be due to the difference in the expression of HypaCas9 protein. We observed a relatively low gene transduction efficiency and low expression level in PC9 cells in comparison to the other two cell lines based on the expression level of EGFP, which was co-expressed with Cas9 via the T2-peptide (Supplementary Fig. 12). Therefore, it is possible that certain levels of Cas9 expression and activity are necessary for the binary induction of genome editing. Another possibility is that different DNA repair pathways are active in different cell types to yield different genome editing outcomes. In PC9 cells, HDR was barely induced even with HypaCas9, which could be because the HDR pathway is not very active in PC9 cells. An investigation of the active DNA repair pathways in these cell lines would be an interesting way to address the observed differences in genome editing outcomes.

We are currently applying the same analytic procedure to other cell types to further address the generality of our findings. Our target cells types include human iPS cells, because we are highly interested in the genome editing outcomes of normal diploid human cells. However, the biggest challenge is the low survival rate of iPS cells, especially in single cell culture^36^; thus, we are optimizing the culture conditions. We can also apply different transduction methods, including lipofection, electroporation of plasmids and RNPs, and viral infection. We will test these various parameters by our SPiS-based strategy to figure out general characteristics of genome editing outcomes in single cells, and also best conditions to induce specific types of genome editing.

Our study reveals the previously unknown but fundamental features of genome editing in single cells including the “binary nature” of genome editing induction. This was only possible with the SPiS. Our findings contribute to the better understanding of the underlying mechanism of induction of genome editing. Moreover, NHEJ often results in gene disruption; thus, the gene functions in a cell may be lost in clones fully edited by NHEJ. This can be very important in genome editing therapy that is dependent on HDR, as by-products of NHEJ might hamper the therapy if the function of a certain gene is completely lost in some cells. Therefore, the analysis of genome editing results in single cells is critical in precise evaluation of therapeutic effects. Our SPiS also provides researchers with a versatile platform to study genome editing in single cells.

## Methods

### Statistical analyses

The transfection and single cell dispensing experiments were performed in triplicate (three biological replicates). Clones isolated by the SPiS that showed no amplification of target sites by PCR or no successful alignments of amplicon sequencing reads by CRISPResso2^27^ were excluded from the analyses. Statistical significance between two groups was assessed by a non-paired two-tailed Student’s *t*-test.

### Plasmids and oligonucleotides for transfection

We generated a vector that co-expresses EGFP and HypaCas9 (pX458-HypaCas9), based on pX458 (Addgene #48138). The single strand DNA donors were all 60 nt and had point mutations in the middle of them. The sequences of the single strand DNA donors and gRNAs that were used in this study are summarized in Supplementary Tables S4 and S5, respectively. All the single strand DNA donors and oligonucleotides for gRNA cloning, which were purified by standard desalting, were purchased from FASMAC, Japan.

### Calculation of the copy number of target genes by karyotyping and CGH analysis

The karyotyping analysis of the cell lines was performed by Nihon Gene Research Laboratories. Karyotypes were analyzed in 10 cells for each cell line. The CGH assays were performed using the SurePrint G3 Human CGH Microarray Kit (Agilent) and the genomic DNA of WTC11 cells as a reference of the diploid cells. The genomic DNA of the cell lines used in this study for the CGH analysis was extracted using the DNeasy Blood and Tissue Kit (QIAGEN) according to the manufacturers’ instructions. The CGH analysis gave peaks of the frequently observed signal intensity, which correspond to the number of chromosomes. Therefore, the chromosomal number most frequently observed in the karyotype analysis was assigned to the most frequent peak in the CGH array (for example, because the most frequently observed chromosomal number was 3 by karyotyping of HEK293T cells, the most frequent peak of the signal intensity in the CGH assay corresponded to 3 copies). Then, we were able to incrementally assign copy numbers to the peaks in the CGH assay around the highest peak to draw a line of fit between the copy number and the CGH peak values. The precise copy number of the target gene was calculated by this line of fit and the signal intensity around the target genes measured by the CGH assay.

### Cell culture, transfection, and cell sorting

HEK293T cells and HeLa cells were maintained in Dulbecco’s modified Eagle medium (DMEM) with high glucose, sodium pyruvate, and L-glutamine (Thermo Fisher Scientific) supplemented with 10% fetal bovine serum (FBS, Thermo Fisher Scientific) and 1% Penicillin-Streptomycin (P/S, Sigma-Aldrich). PC9 cells were maintained in Roswell Park Memorial Institute (RPMI) medium with L-glutamine (Nacalai Tesque) supplemented with 10% FBS (Thermo Fisher Scientific) and 1% P/S. For transfection, 2×10^5^ cells were plated in a well of a 12-well plate coated with 80 μg/ml Growth Factor Reduced Matrigel Basement Membrane Matrix (BD Biosciences). One day later, the cells were transfected with 720 ng of pX458 or pX458-HypaCas9 with a gRNA targeting RBM20, ATP7B, or GRN, and 80 ng of oligonucleotide donor DNA using 2.4 μl of Lipofectamine 2000 (Thermo Fisher Scientific) for HEK293T cells and HeLa cells, and 2.0 μl of Lipofectamine 3000 (Thermo Fisher Scientific) for PC9 cells, respectively, per well, according to the manufacturers’ instructions. After 24 hours, cells were detached by 0.25% Trypsin-EDTA (Thermo Fisher Scientific) and EGFP-positive cells were sorted using On-ship Sort (On-chip Biotechnologies). The isolated EGFP-positive cells were dispensed one by one into Matrigel-coated 96-well plates using the On-chip Single Particle isolation System (SPiS) (Fig. 2c). Subsequently, 100 μl/well of conditioned medium was added to the plates with dispended cells.

One week after single cell dispensing by SPiS, the medium was changed. Two weeks after dispensing, for HEK293T cells and HeLa cells, the surviving colonies were first detached by Trypsin-EDTA, then the cells were resuspended in DMEM supplemented with 20% FBS and 1% P/S to evenly re-distribute the cells within the wells of a 96-well plate. The cells were cultured in this media until they reached confluence.

### Preparation of multiplexed amplicon sequencing library

Genomic DNA was extracted from the cells in 96-well plates as described previously^31^. The DNA was resuspended in 30 μl/well of water. Targeted sites were amplified by the first PCR round using primers with homology to the region of interest and the Illumina forward and reverse adapters (Supplementary Table 6). The first PCR round consisted of 0.3 μl each of 100 μM forward and reverse primer, 1.0 μl of genomic DNA, 2.0 μl of 2 mM dNTPs, 5.0 μl of 2× PCR Buffer KOD FX (Toyobo), 0.1 μl of KOD FX enzyme (Toyobo), 2 μl of betaine (Fuji Film), and 0.3 μl of water. The thermal cycling settings were as follows: 95 °C for 2 min, then 30 cycles of (95 °C for 30 sec, 60 °C for 30 sec, and 72 °C for 1 min), followed by a final 72 °C extension for 3 min. The first PCR products were diluted by adding 90 μl of water. Then, the DNA barcodes and illumina adaptors were added to the amplicons in the second PCR round, which consisted of 0.3 μl each of 100 μM unique forward and reverse index barcoding primers^37^ (Supplementary table 7), 0.5 μl of the diluted first PCR product, 2 μl of 2 mM dNTPs, 5.0 μl of 2× PCR Buffer KOD FX, 0.1 μl of KOD FX enzyme, and 1.8 μl of water. The thermal cycling settings were as follows: 98°C for 3 min, then 30 cycles of (98 °C for 30 sec, 57 °C for 30 sec, and 72 °C for 1 min), followed by a final 72 °C extension for 3 min. The PCR products were electrophoresed in 2% agarose gel. NucleoSpin Gel and PCR Clean-up Midi kit (MACHEREY-NAGEL) were used to extract the pooled PCR products from 24 samples in 200 μl of water. The DNA concentrations of these library mixtures of 24 samples were quantified using the GenNext NGS Library Quantification Kit (Toyobo). All libraries were then mixed in equimolar amounts, 20% PhiX Control v3 (illumina) was added for amplicon sequencing. Sequencing was performed with MiSeq (Illumina) using the Miseq v2 reagent kit (Illumina) or Miseq Reagent kit v2 Nano (illumina) according to the manufacturer’s instructions.

### Amplicon sequencing data analysis

Fastq files generated by MiSeq were imported into the CLC Genomics Workbench 21 (QIAGEN). Adapter sequences were removed and demultiplexed using the DNA Index. The data were then analyzed by CRISPResso2^27^ (https://github.com/pinellolab/CRISPResso2) in the CRISPResso Batch mode. CRISPResso2 was installed as recommended using a Docker containerization system. The commands for the CRISPResso2 analysis used in this study are listed in Supplementary Table 8. Alleles with ≤5% frequency were excluded from the analysis as they were expected to be either sequence errors or very minor populations generated during cell proliferation from single cells. In the CRISPResso2 analysis, any alleles with deletions spanning the nucleotides for single nucleotide substitutions were characterized as “ambiguous”. Therefore, we classified these “ambiguous” alleles as NHEJ alleles in this study.

### Creating a model in which alleles were evenly distributed

Models assuming that the WT, NHEJ, HDR, and HDR+NHEJ alleles were randomly induced in all target alleles were built by distributing these edited alleles at their observed overall frequencies. The number of target alleles per cell was calculated by karyotyping and a CGH analysis, as described above. For example, when ATP7B was targeted in HEK293T cells, one cell has three ATP7B alleles. If the overall frequencies of WT, NHEJ, HDR, and HDR+NHEJ were 40%, 30%, 20%, and 10%, respectively, and 100 HEK293T cell clones were isolated, the expected number of clones with WT, NHEJ, and HDR alleles would be two (100×(0.4×0.3×0.2)=2.4, rounded up to 2). We calculated the expected clone numbers of all the possible combinations of WT, NHEJ, HDR, and HDR+NHEJ alleles, and combined them to build the models.

## Data availability

Raw data of multiplexed amplicon sequencing in this study are available in the DDBJ Sequence Read Archive under the accession numbers DRA013570.

## Acknowledgements

We thank Drs. Y. Fujimura and K. Takeda (On-chip Biotechnologies) for their helpful discussions.

This work was supported by JSPS Grant-in-Aid for Challenging Research (Pioneering) 20K21409, Grant-in-Aid for Scientific Research (B) 20H03442, Interstellar Initiative from AMED (20jm0610032h0001), Takeda Science Foundation Medical Research Grant, Sumitomo Foundation Grant For Basic Science Research Projects, Ichiro Kanehara Foundation Grant to Y.M., and a JSPS Grant-in-Aid for Early-Career Scientists (18K15054) to G.T.

## Author contributions

G.T. and Y.Miyaoka conceived the idea of this study and designed the experiments. G.T., D.K., and M.M. isolated genome edited HEK293T cell clones with technical help from Y.Morishita.

D.K. and M.M. prepared libraries for amplicon sequencing. G.T., D.K., M.M., and Y.Miyaoka analyzed the results. Y.Miyaoka supervised the project. G.T. and Y.Miyaoka wrote the manuscript with help from all of the authors.

