## Supplementary Figures and Tables for "Genome Editing Is Induced in a Binary Manner in Single Human Cells"

### Supplementary Figure. 1

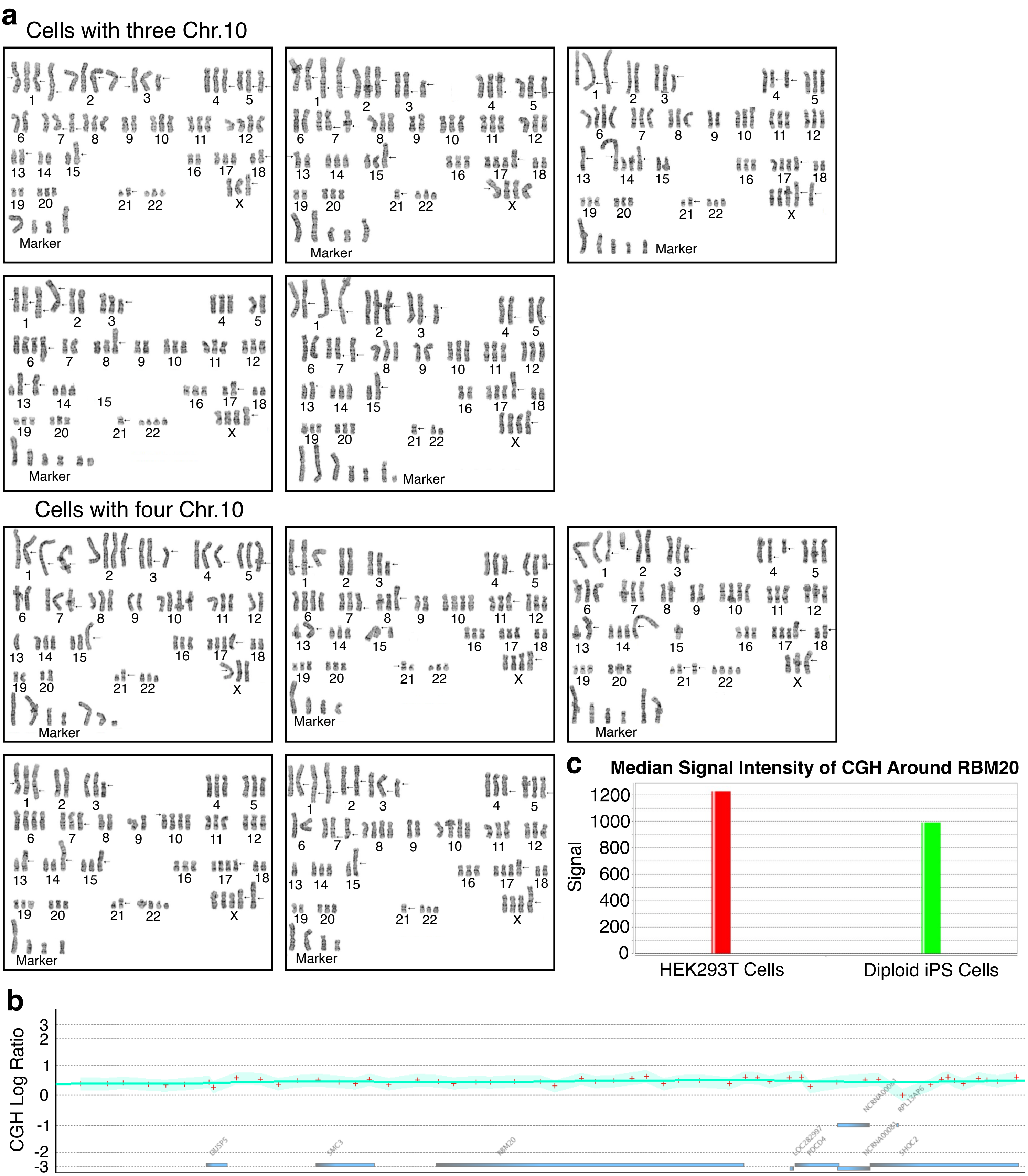

**Supplementary Figure 1. Karyotyping and a CGH analysis of HEK293T cells to estimate the copy number of RBM20.**

(a) Karyotypes of ten HEK293T cells. Five cells had three chromosome 10s, and the other five had four chromosome 10s. Please note that two of them are also shown in Fig. 1b. (b) Scattered plot of the relative CGH signal of HEK293T cells in comparison to diploid human iPS cells around the RBM20 gene. The relative CGH signals of HEK293T cells normalized by the CGH signals of iPS cells are represented by +. This panel shows a wider genomic region than Fig. 1f. There were no detectable microduplications or microdeletions around the RBM20 locus. (c) The median signal intensity of CGH around RBM20. The median signal intensities of HEK293T cells and iPS cells in the genomic region shown in (b). By applying these values to calculate the CGH log ratio and applying it to the formula shown in Fig. 1e, the copy number of RBM20 was calculated to be 3.54.

#### Supplementary Figure. 2

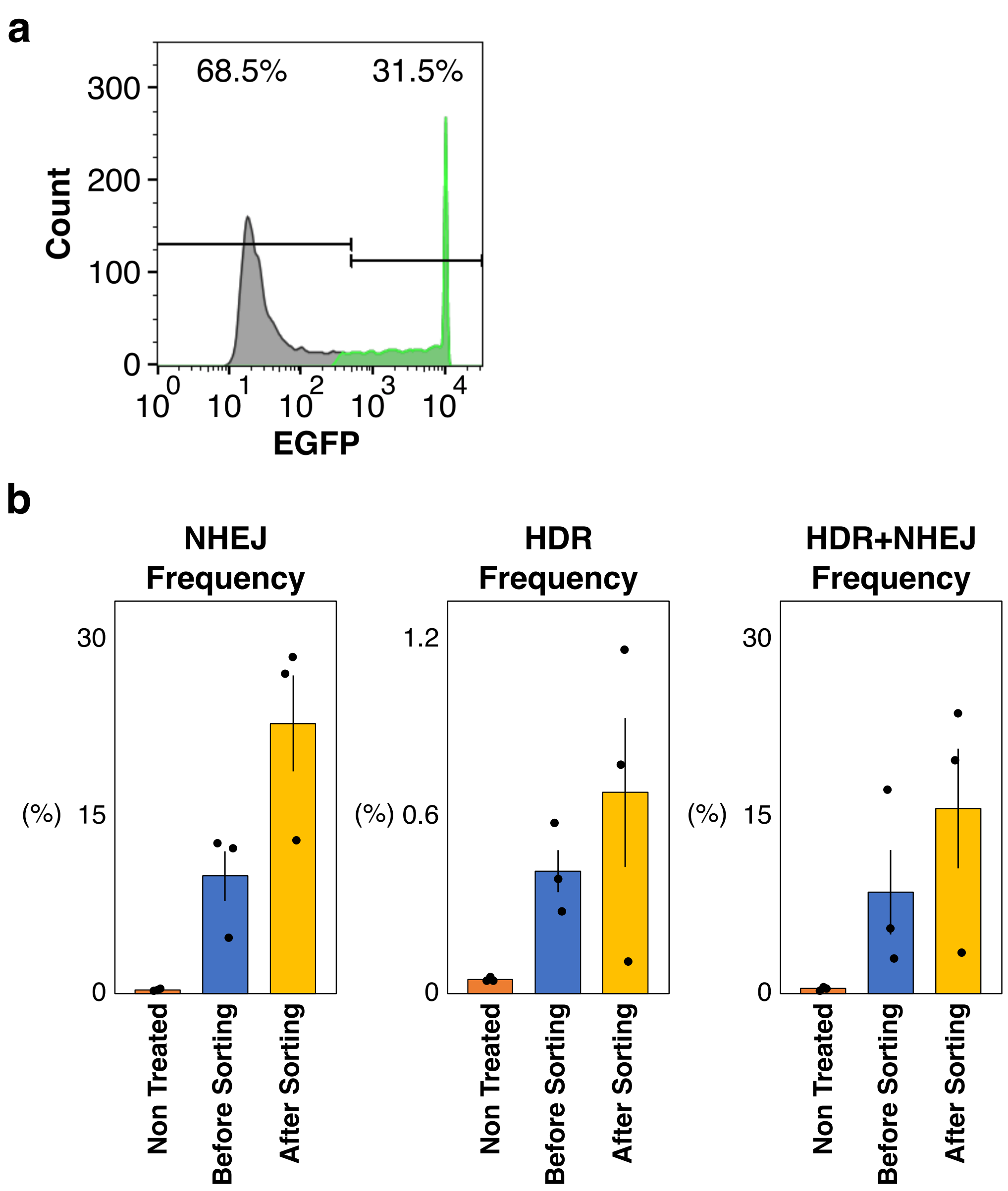


**Supplementary Figure 2. Sorting of EGFP-positive HEK293T cells.**

(a) Histogram showing the sorting gate of EGFP-positive HEK293T cells. (b) Frequencies of NHEJ, HDR, and HDR+NHEJ alleles before and after cell sorting. Cas9, gRNA targeting RBM20, and EGFP were co-expressed in HEK293T cells and EGFP-positive cells were sorted. The frequencies of NHEJ, HDR, and HDR+NHEJ alleles were analyzed by amplicon sequencing. Values ± S.E. are shown (n=3).

#### Supplementary Figure. 3

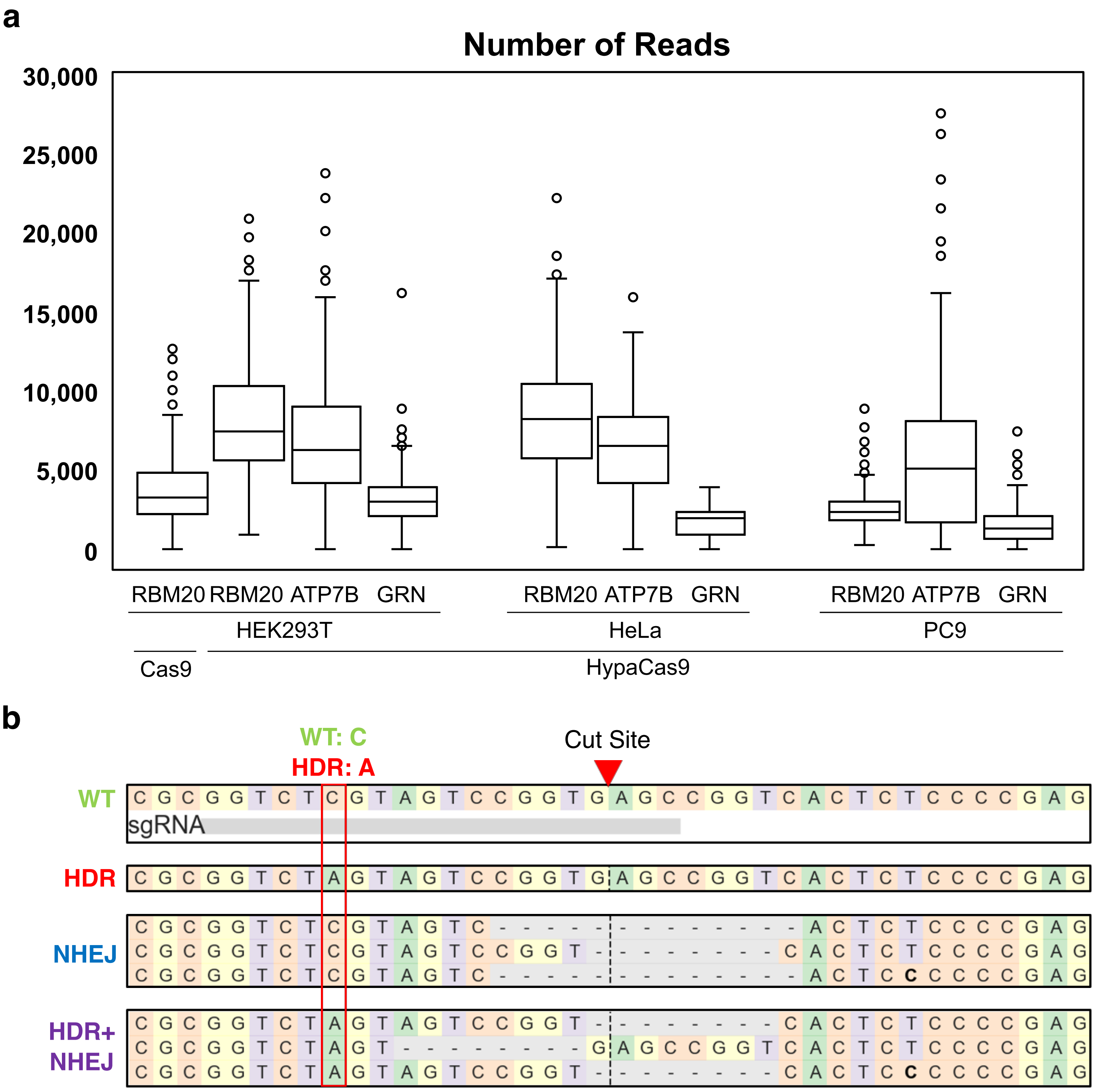


**Supplementary Figure 3. Amplicon sequencing of isolated clones.**

(a) The number of reads sequenced by amplicon sequencing in this study. (b) Example reads of amplicon sequencing of RBM20 targeting in HEK293T cells. The cut site and mutated site are shown.

### Supplementary Figure. 4

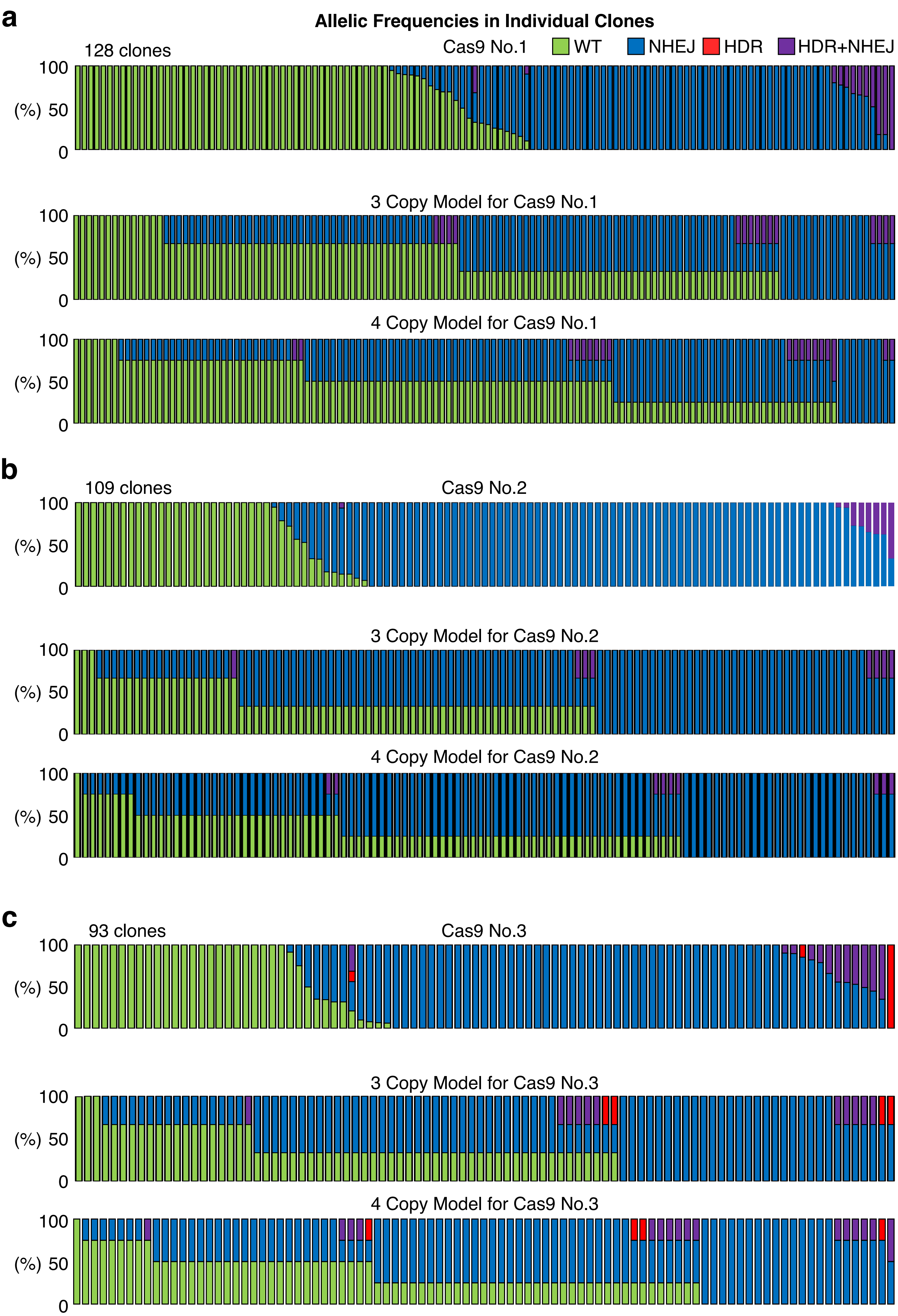

**Supplementary Figure 4. Models assuming random RBM20 R636S mutagenesis by Cas9 in HEK293T cells.**

(a-c) Models of the distribution of HEK293T cell clones with different genome editing outcomes by Cas9, if genome editing randomly occurred at the overall frequencies in HEK293T cells with three or four copies of RBM20 in experiment No. 1 (a), No. 2 (b), and No. 3 (c). The top row in each panel shows the observed distribution of the isolated HEK293T cell clones. Note that the original data and the 3 and 4 copy models for Cas9 No. 3 are also shown in Fig. 3.

### Supplementary Figure. 5

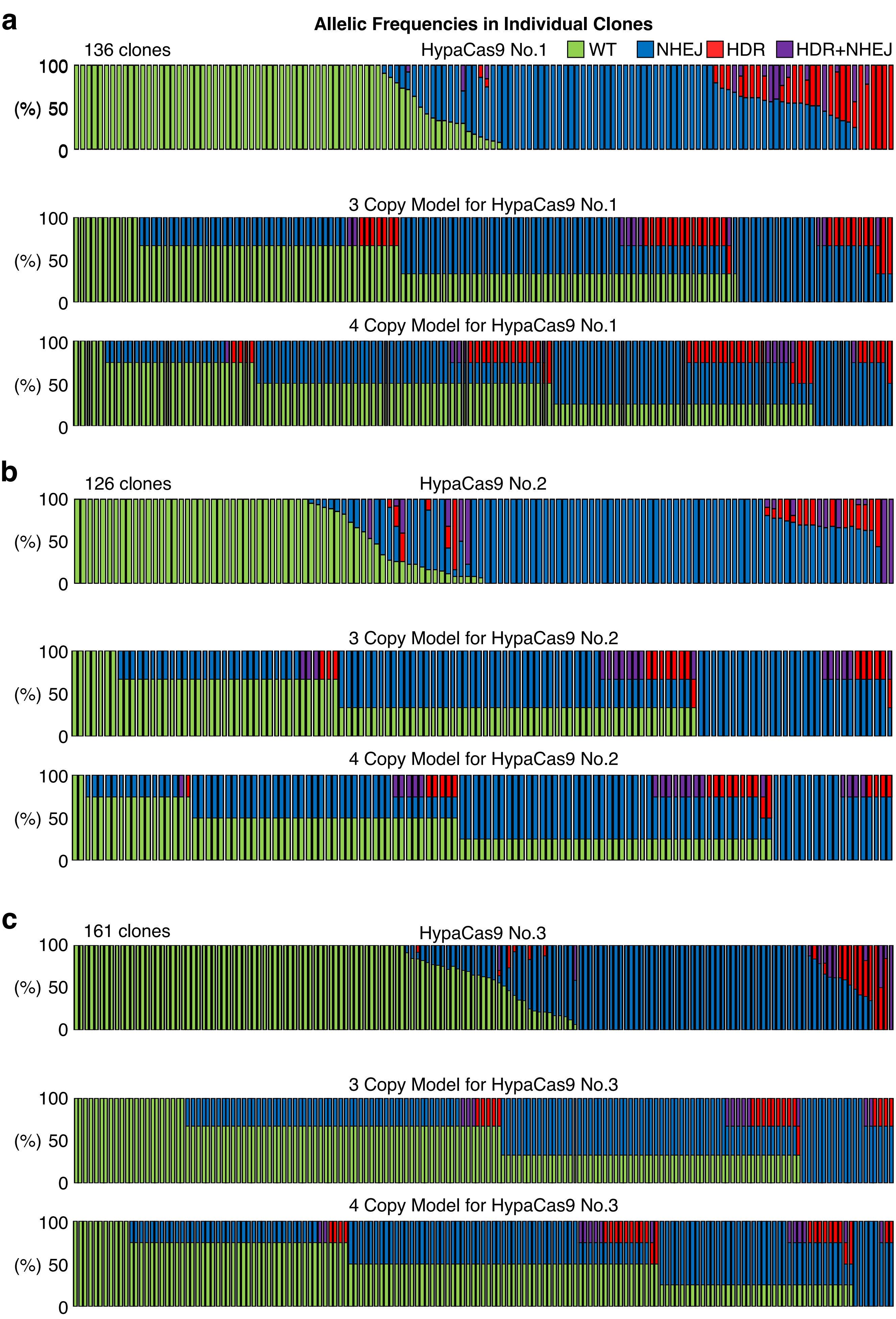

**Supplementary Figure 5. Models assuming random RBM20 R636S mutagenesis by HypaCas9 in HEK293T cells.**

(a-c) Models of the distribution of HEK293T cell clones with different genome editing outcomes by HypaCas9, if genome editing randomly occurred at the overall frequencies in HEK293T cells with three or four copies of RBM20 in experiment No. 1 (a), No. 2 (b), and No. 3 (c). The top row in each panel shows the observed distribution of the isolated HEK293T cell clones. Note that the original data and the 3 and 4 copy models for HypaCas9 No. 3 are also shown in Fig. 4.

### Supplemental Figure 6

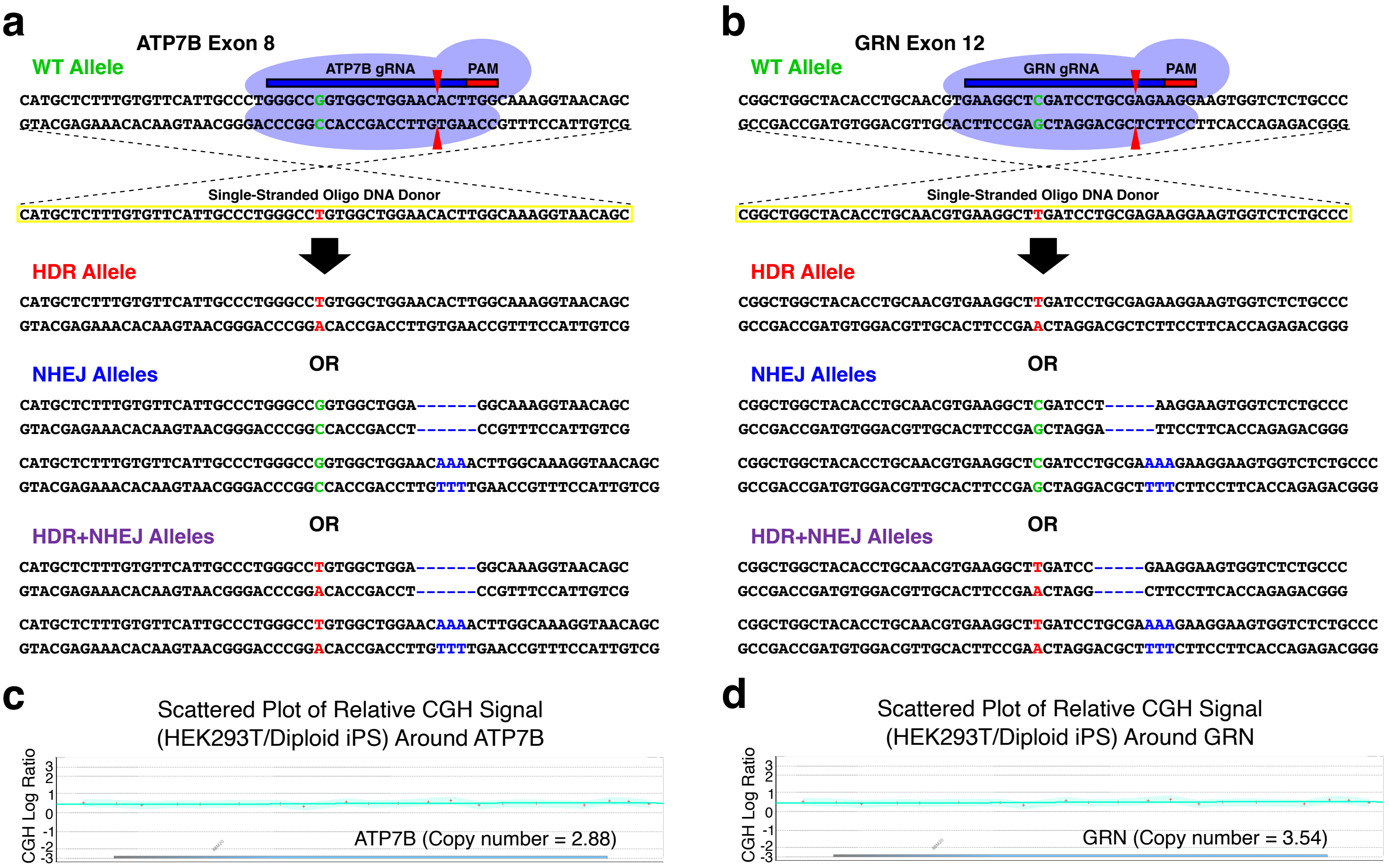


**Supplementary Figure 6. Design of ATP7B R778L and GRN R493X targeting, and estimation of copy numbers of ATP7B and GRN in HEK293T cells.**

(a, b) Design of genome editing in ATP7B (a) and GRN (b), respectively. We introduced the R778L (c.2333C>G) (a) and R493X (c.1477C>T) (b) mutations using HypaCas9 and a single-stranded oligonucleotide donor DNA. The resulting HDR alleles have a single nucleotide substitution from C to G (a) and C to T (b), whereas the NHEJ alleles have various insertions and deletions. (c, d) Scattered plots of the relative CGH signal of HEK293T cells in comparison to diploid human iPS cells around the ATP7B (c) and GRN (d) genes, respectively. The relative CGH signals of HEK293T cells in comparison to diploid iPS cells are represented by +. No microduplications or microdeletions were detected around the ATP7B (c) or GRN (d) loci. The median CGH log ratios in these genomic regions shown here were applied to the formula shown in Fig. 1e to calculate the copy numbers of ATP7B (c) and GRN (d) were 2.88 (c) and 3.54 (d), respectively.

### Supplementary Figure. 7

**a**

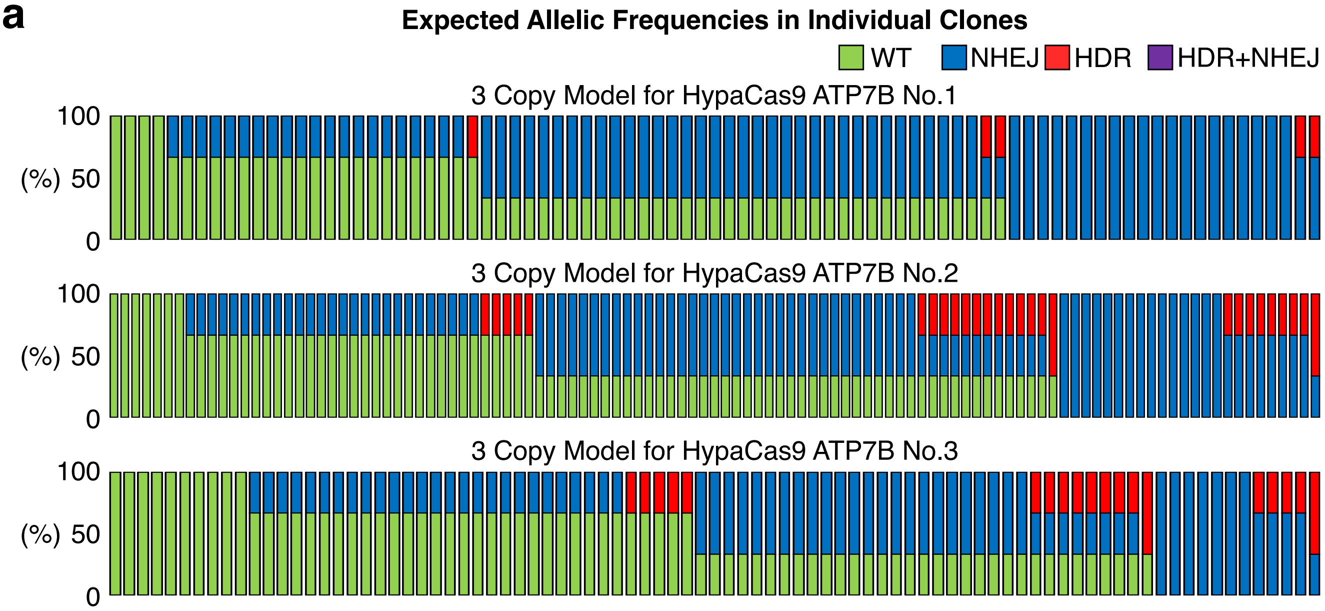


**b**

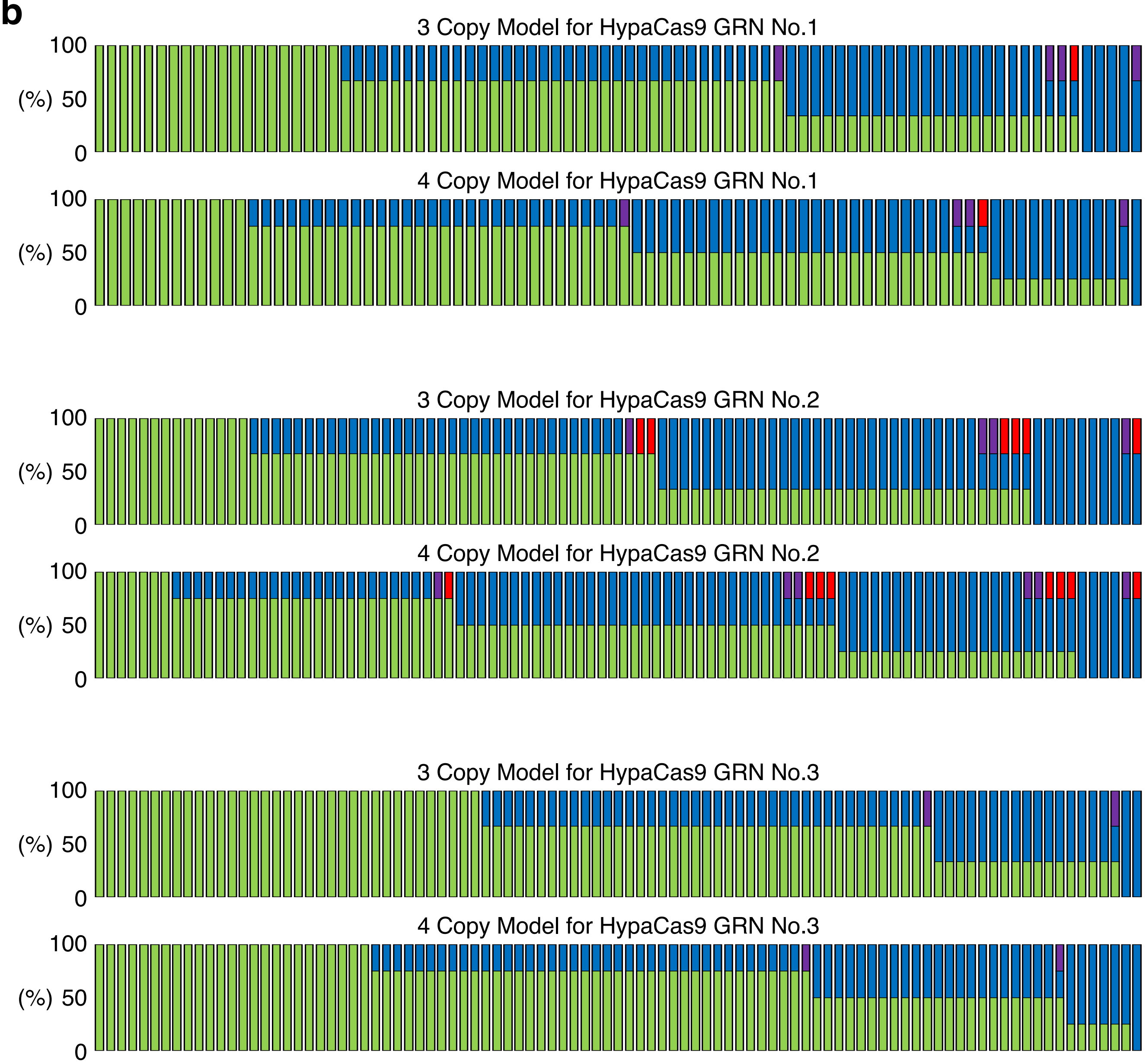

**Supplementary Figure 7. Models assuming random ATP7B R778L and GRN R493X mutagenesis by HypaCas9 in HEK293T cells.**

(a-b) Models of the distribution of HEK293T cell clones with different genome editing outcomes by HypaCas9, if genome editing randomly occurred at the overall frequencies in HEK293T cells with three copies of ATP7B (a) and three or four copies of GRN (b) in experiment Nos. 1-3.

### Supplementary Figure. 8

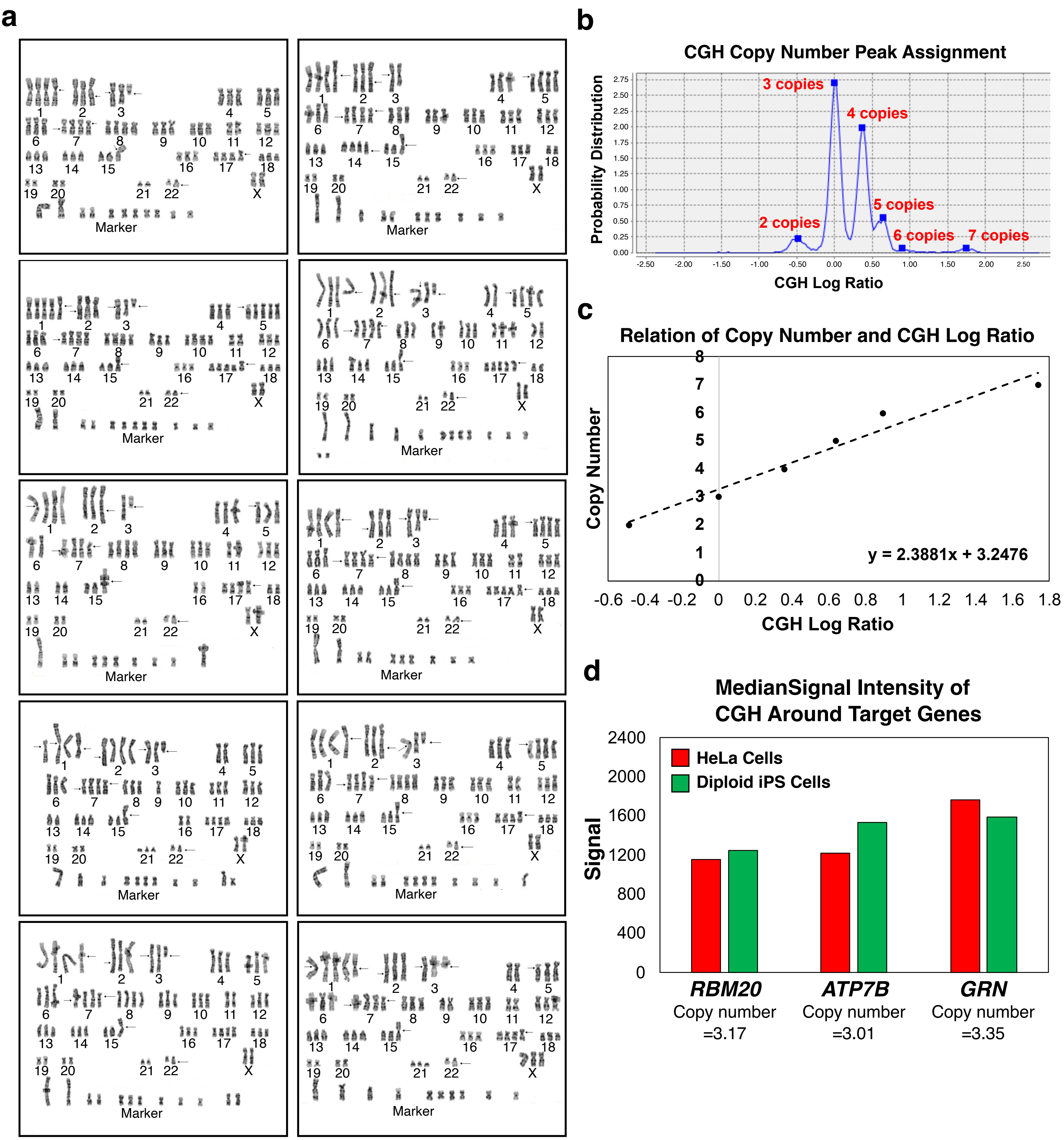

**Supplementary Figure 8. Karyotyping and a CGH analysis to estimate the copy number of RBM20, ATP7B, and GRN in HeLa cells.**

(a) Karyotypes of ten HeLa cells. (b) CGH copy number peak assignment. In the CGH analysis, there were several peaks of the CGH signal ratio between HeLa cells and diploid iPS cells based on the chromosomal numbers. The highest peak corresponded to three copies per cell, and the other peaks were incrementally assigned to the other copy numbers. (c) Line of fit between the copy number and the CGH log ratio based on the peak assignment shown in (b). (d) The median signal intensities of CGH around the RBM20, ATP7B, and GRN genes. By applying these values to calculate the CGH log ratios, which were then applied to the formula shown in (c), the copy numbers of RBM20, ATP7B, and GRN, in HeLa cells were 3.17, 3.01, and 3.35, respectively.

### Supplementary Figure. 9

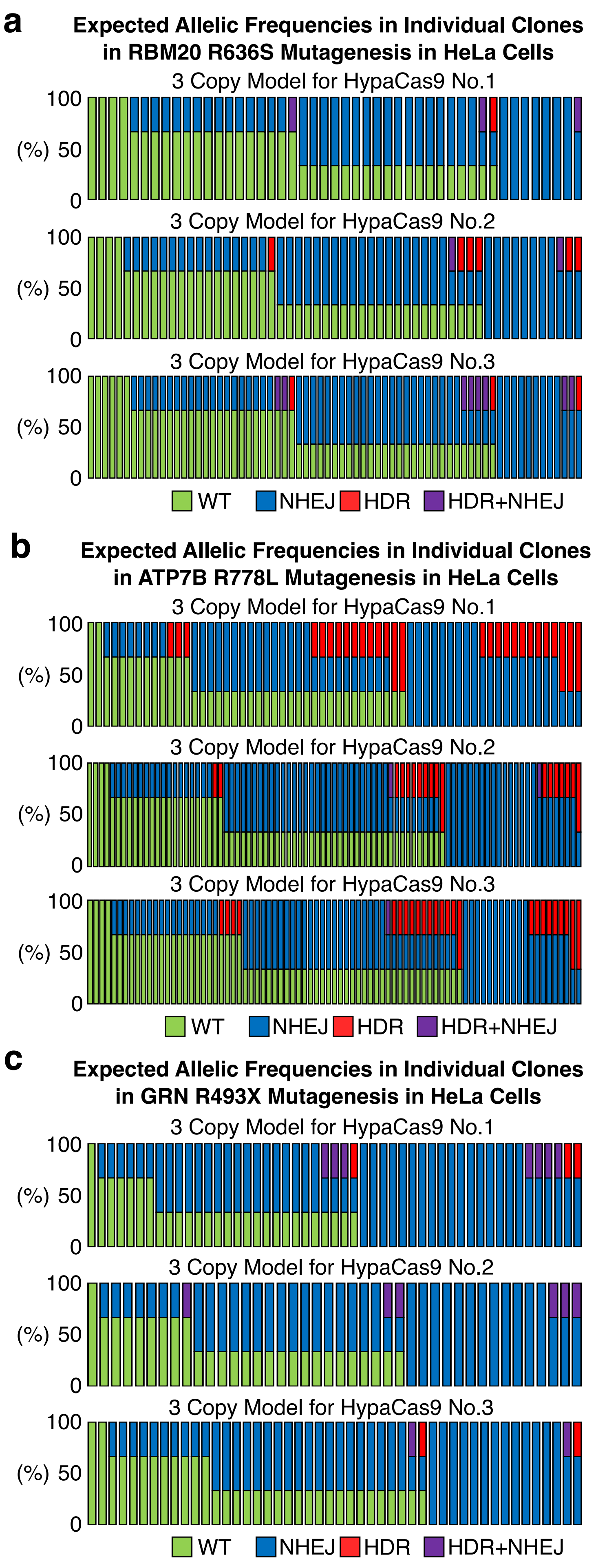


**Supplementary Figure 9. Models assuming random RBM20 R636S, ATP7B R778L, and GRN R493X mutagenesis by HypaCas9 in HeLa cells.**

(a-c) Models of the distribution of HeLa cell clones with different genome editing outcomes by HypaCas9, if genome editing randomly occurred at the overall frequencies in HeLa cells with three copies of RBM20 (a), ATP7B (b), and GRN (c) in experiment Nos. 1-3.

### Supplementary Figure. 10

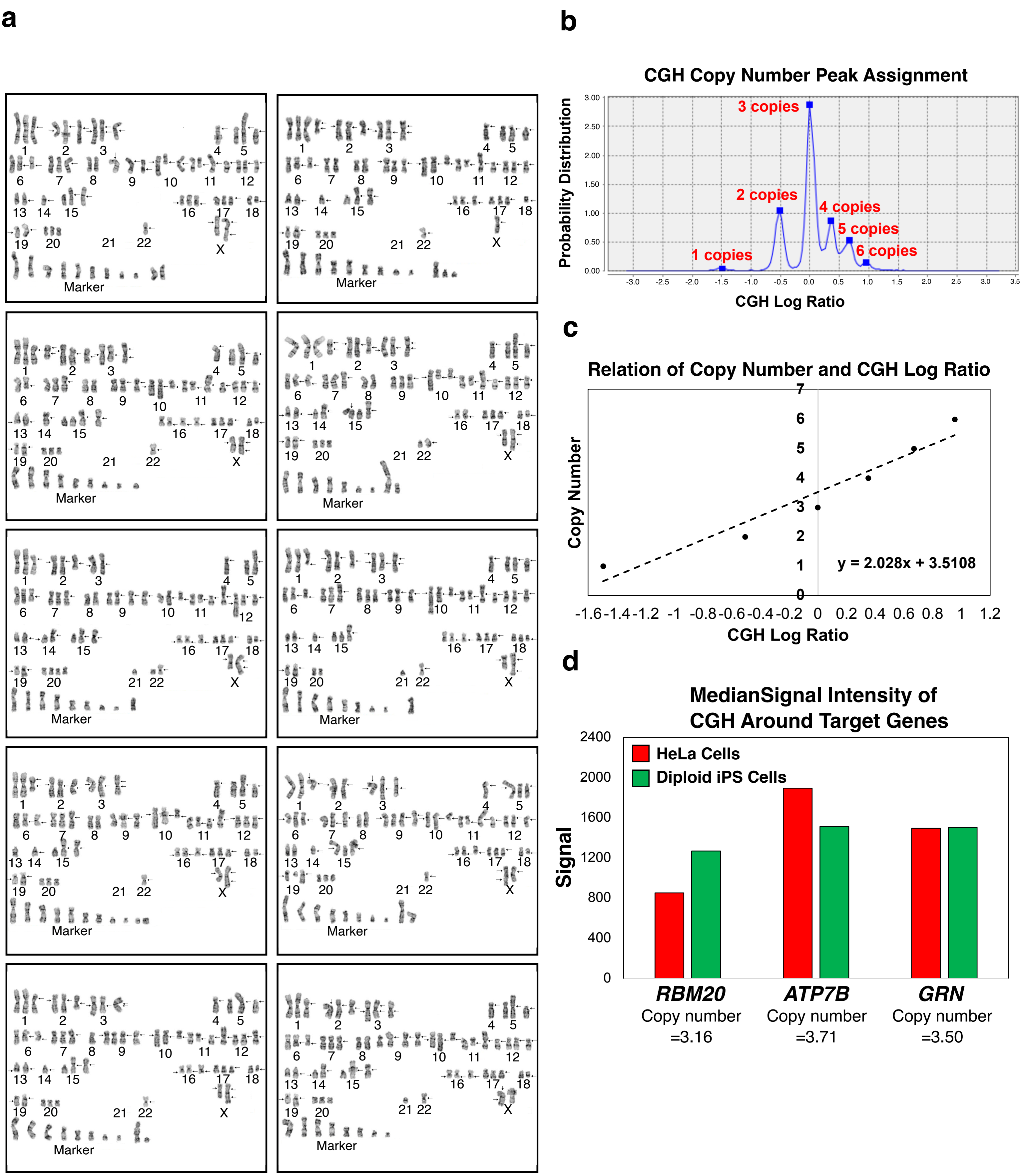

**Supplementary Figure 10. Karyotyping and a CGH analysis to estimate the copy number of RBM20, ATP7B, and GRN in PC9 cells.**

(a) Karyotypes of ten PC9 cells. (b) CGH copy number peak assignment. In the CGH analysis, there were several peaks of the CGH signal ratio between PC9 cells and diploid iPS cells based on the chromosomal numbers. The highest peak corresponded to three copies per cell, and the other peaks were incrementally assigned to the other copy numbers. (c) Line of fit between the copy number and the CGH log ratio based on the peak assignment shown in (b). (d) The median signal intensities of CGH around the RBM20, ATP7B, and GRN genes. By applying these values to calculate the CGH log ratios, which were then applied to the formula shown in (c), the copy numbers of RBM20, ATP7B, and GRN in PC9 cells were 3.16, 3.71, and 3.50, respectively.

### Supplementary Figure. 11

a

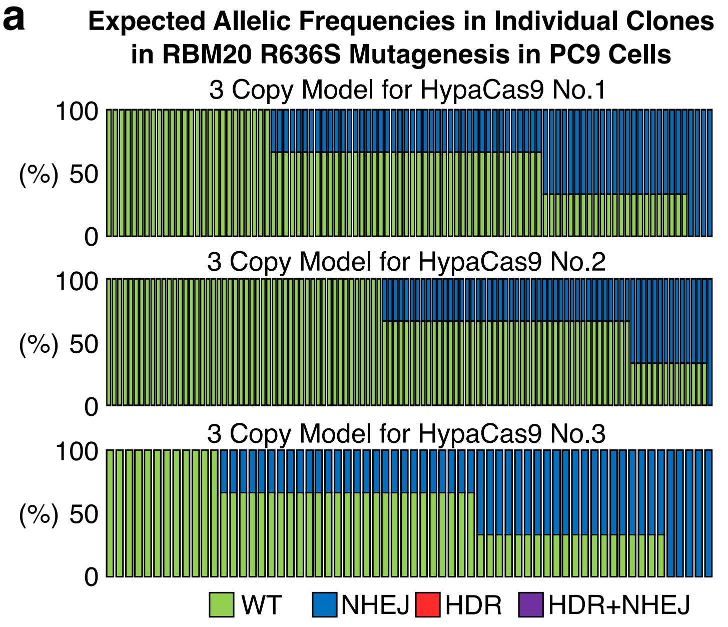


b

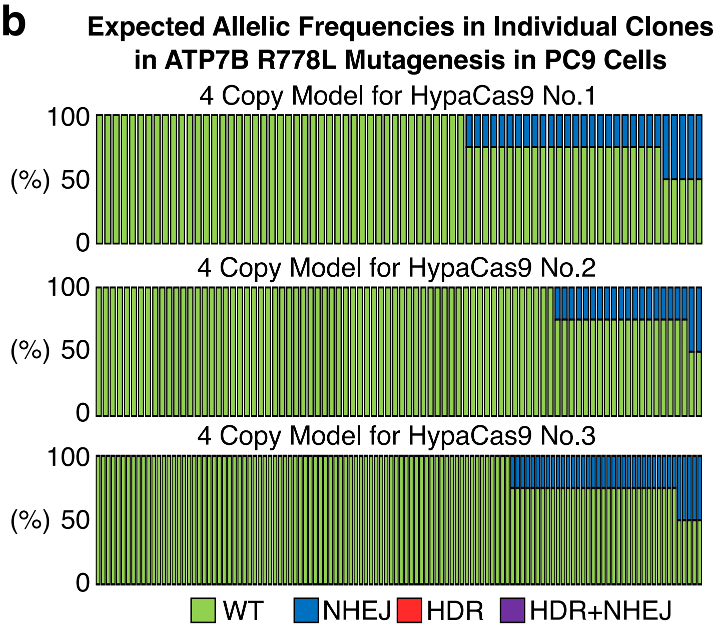


**Supplementary Figure 11. Models assuming random RBM20 R636S, ATP7B R778L, and GRN R493X mutagenesis by HypaCas9 in PC9 cells.**

(a-c) Models of the distribution of PC9 cell clones with different genome editing outcomes by HypaCas9, if genome editing randomly occurred at the overall frequencies in PC9 cells with three copies of RBM20 (a), four copies of ATP7B (b), and three or four copies of GRN (c) in experiment Nos. 1-3.

c

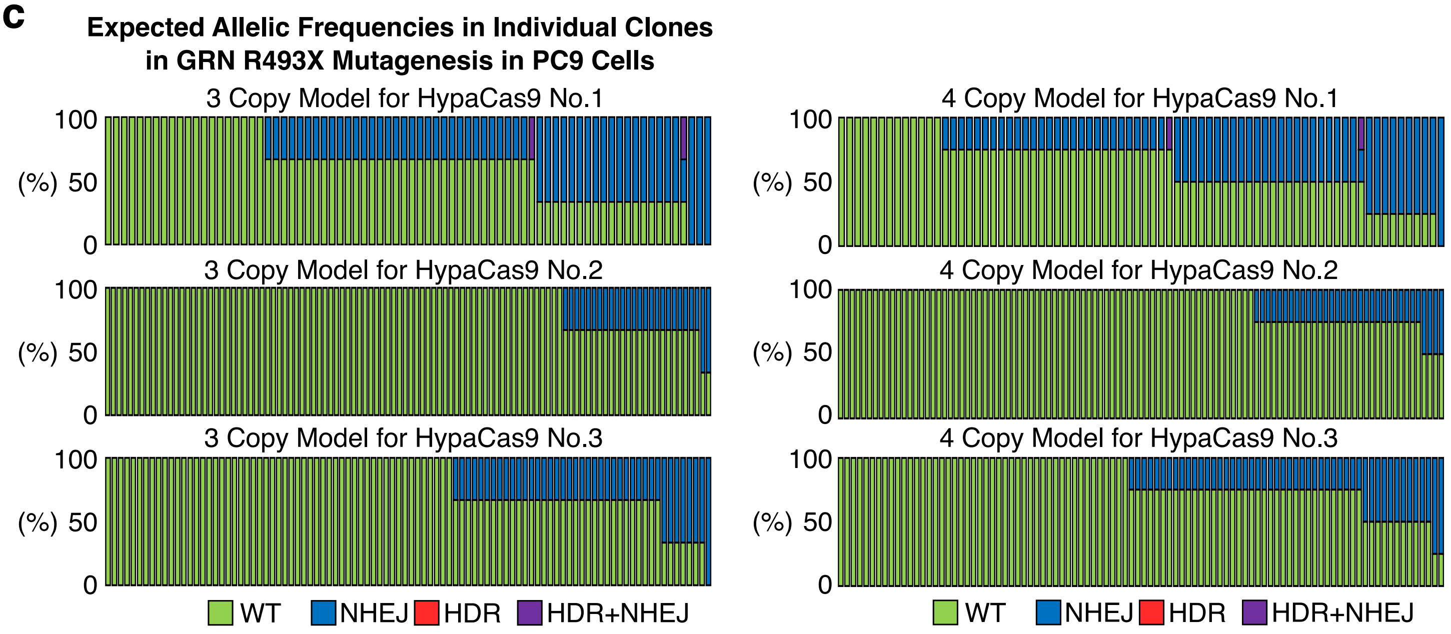

#### Supplementary Figure. 12

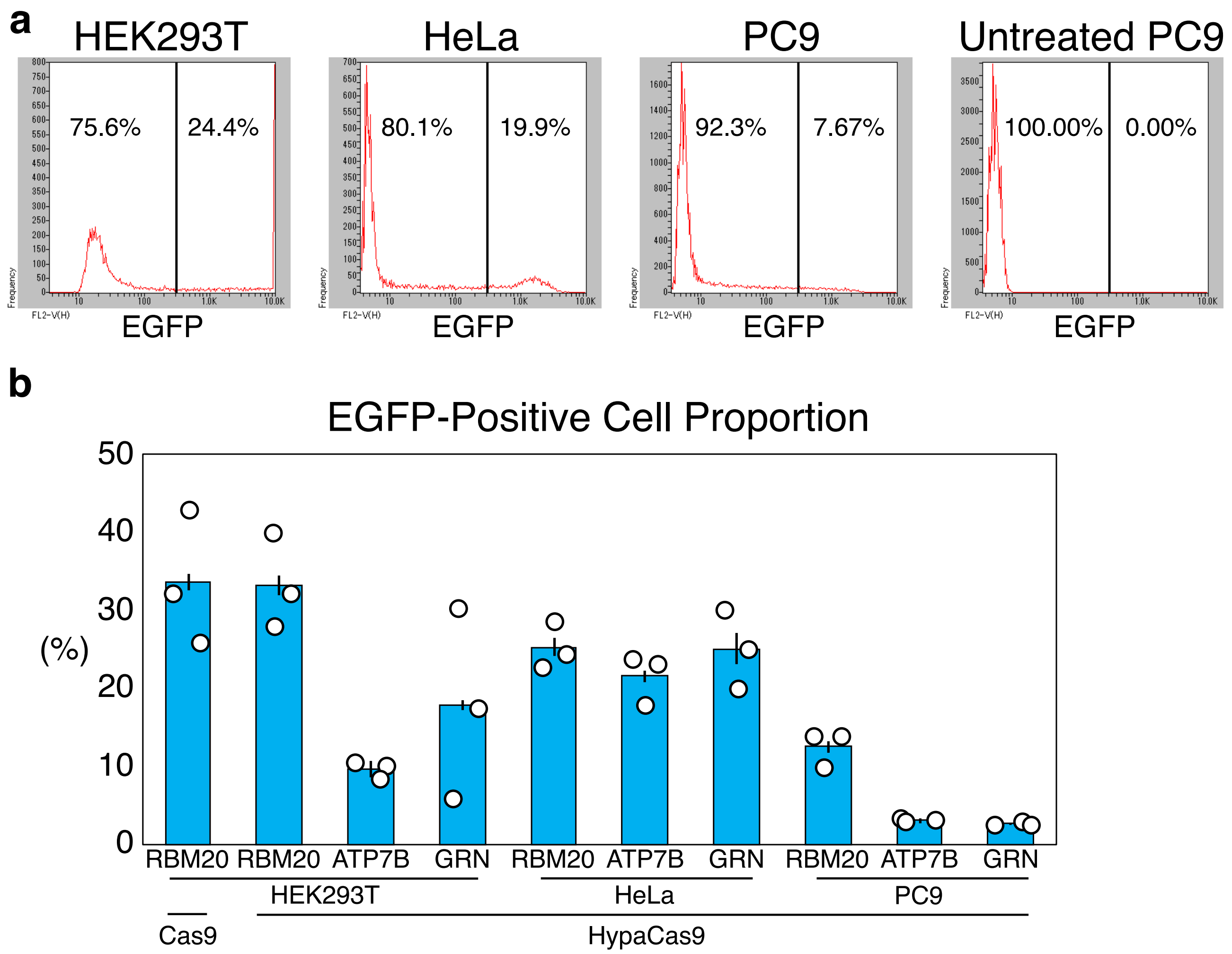


**Supplementary Figure 12. EGFP-positive populations after co-expression of Cas9/HypaCas9, gRNA, and EGFP in HEK293T cells, HeLa cells, and PC9 cells.**

(a) Representative histograms of EGFP-positive populations in HEK293T cells, HeLa cells, and PC9 cells 2 days after transfection of the plasmids to target RBM20. PC9 cells without any treatment are shown as a negative cell population. The expression level of EGFP in HEK293T cells was saturated in this histogram. (b) EGFP-positive cell populations determined by flow cytometry. Values ± S.E. are shown (n=3).

**Supplementary Table 1. Chromosomal numbers in HEK293T cells**

We counted the chromosomal numbers in ten HEK293T cells. There were 1, 8, 64, 118, 37, and 2 cases where the chromosomal number was 0, 1, 2, 3, 4, and 5, respectively. The most frequent chromosomal number was three. We observed no Y chromosomes, as HEK293T cells were derived from a female. There were no cases where the chromosomal number was more than 6.

| # | Cell 1 | Cell 2 | Cell 3 | Cell 4 | Cell 5 | Cell 6 | Cell 7 | Cell 8 | Cell 9 | Cell 10 |
|---|---|---|---|---|---|---|---|---|---|---|
| 1 | 4 | 3 | 3 | 4 | 3 | 4 | 4 | 4 | 4 | 4 |
| 2 | 2 | 2 | 2 | 2 | 2 | 3 | 3 | 2 | 4 | 4 |
| 3 | 3 | 3 | 3 | 3 | 3 | 3 | 3 | 3 | 3 | 3 |
| 4 | 3 | 3 | 3 | 3 | 3 | 2 | 3 | 3 | 3 | 3 |
| 5 | 3 | 3 | 3 | 2 | 3 | 2 | 3 | 3 | 3 | 3 |
| 6 | 3 | 4 | 4 | 4 | 4 | 2 | 2 | 2 | 2 | 2 |
| 7 | 3 | 3 | 3 | 2 | 3 | 3 | 3 | 3 | 3 | 3 |
| 8 | 2 | 3 | 3 | 3 | 2 | 3 | 3 | 4 | 3 | 3 |
| 9 | 2 | 2 | 2 | 2 | 2 | 2 | 2 | 2 | 2 | 2 |
| 10 | 4 | 3 | 4 | 3 | 4 | 3 | 3 | 4 | 4 | 3 |
| 11 | 3 | 3 | 3 | 3 | 3 | 3 | 3 | 3 | 3 | 3 |
| 12 | 3 | 3 | 3 | 3 | 3 | 3 | 3 | 3 | 2 | 4 |
| 13 | 2 | 1 | 2 | 3 | 2 | 2 | 2 | 1 | 1 | 2 |
| 14 | 4 | 4 | 3 | 3 | 3 | 3 | 3 | 3 | 3 | 2 |
| 15 | 1 | 2 | 2 | 0 | 3 | 2 | 3 | 2 | 3 | 2 |
| 16 | 3 | 3 | 3 | 3 | 3 | 2 | 3 | 3 | 3 | 2 |
| 17 | 4 | 4 | 3 | 2 | 4 | 4 | 4 | 4 | 4 | 3 |
| 18 | 2 | 2 | 2 | 2 | 2 | 2 | 2 | 2 | 2 | 2 |
| 19 | 3 | 3 | 3 | 3 | 3 | 3 | 2 | 2 | 2 | 2 |
| 20 | 3 | 3 | 3 | 3 | 3 | 3 | 3 | 3 | 2 | 3 |
| 21 | 3 | 2 | 2 | 1 | 2 | 1 | 1 | 1 | 2 | 2 |
| 22 | 4 | 3 | 3 | 4 | 4 | 2 | 3 | 3 | 3 | 3 |
| X | 3 | 5 | 4 | 4 | 5 | 4 | 4 | 4 | 3 | 3 |
| Y | 0 | 0 | 0 | 0 | 0 | 0 | 0 | 0 | 0 | 0 |

**Supplementary Table 2. Chromosomal numbers in HeLa cells**

We counted the chromosomal numbers in ten HeLa cells. There were 1, 72, 114, 37, and 6 cases where the chromosomal number was 1, 2, 3, 4, and 5, respectively. The most frequent chromosomal number was three. We observed no Y chromosomes, as HeLa cells were derived from a female. There were no cases where the chromosomal number was 0 or more than 6.

| # | Cell 1 | Cell 2 | Cell 3 | Cell 4 | Cell 5 | Cell 6 | Cell 7 | Cell 8 | Cell 9 | Cell 10 |
|---|---|---|---|---|---|---|---|---|---|---|
| 1 | 4 | 5 | 4 | 4 | 4 | 4 | 4 | 4 | 4 | 5 |
| 2 | 3 | 3 | 3 | 4 | 3 | 3 | 3 | 3 | 2 | 3 |
| 3 | 3 | 3 | 2 | 3 | 3 | 2 | 3 | 3 | 3 | 3 |
| 4 | 3 | 3 | 3 | 2 | 3 | 3 | 2 | 3 | 3 | 2 |
| 5 | 3 | 5 | 3 | 3 | 3 | 4 | 4 | 4 | 4 | 3 |
| 6 | 3 | 3 | 2 | 3 | 3 | 3 | 3 | 3 | 3 | 3 |
| 7 | 4 | 4 | 4 | 4 | 4 | 4 | 4 | 4 | 4 | 4 |
| 8 | 4 | 4 | 4 | 3 | 4 | 3 | 3 | 4 | 4 | 3 |
| 9 | 3 | 3 | 3 | 1 | 3 | 3 | 2 | 3 | 3 | 2 |
| 10 | 3 | 4 | 3 | 3 | 3 | 3 | 3 | 3 | 3 | 3 |
| 11 | 2 | 2 | 2 | 3 | 2 | 3 | 3 | 2 | 2 | 2 |
| 12 | 3 | 3 | 3 | 3 | 3 | 3 | 2 | 3 | 3 | 3 |
| 13 | 3 | 3 | 2 | 2 | 3 | 3 | 3 | 3 | 3 | 3 |
| 14 | 3 | 3 | 2 | 3 | 3 | 4 | 3 | 3 | 3 | 3 |
| 15 | 3 | 3 | 3 | 3 | 3 | 3 | 3 | 3 | 3 | 3 |
| 16 | 3 | 3 | 2 | 2 | 3 | 3 | 3 | 3 | 3 | 3 |
| 17 | 4 | 5 | 4 | 4 | 4 | 3 | 5 | 5 | 4 | 4 |
| 18 | 3 | 3 | 2 | 3 | 3 | 2 | 2 | 3 | 3 | 3 |
| 19 | 2 | 2 | 2 | 2 | 2 | 2 | 2 | 2 | 2 | 2 |
| 20 | 2 | 2 | 2 | 2 | 2 | 2 | 2 | 2 | 2 | 2 |
| 21 | 2 | 2 | 2 | 3 | 2 | 2 | 2 | 2 | 2 | 3 |
| 22 | 2 | 2 | 2 | 2 | 2 | 2 | 2 | 2 | 2 | 2 |
| X | 2 | 2 | 2 | 2 | 2 | 2 | 2 | 2 | 2 | 3 |
| Y | 0 | 0 | 0 | 0 | 0 | 0 | 0 | 0 | 0 | 0 |

**Supplementary Table 3. Chromosomal numbers in PC9 cells**

We counted the chromosomal numbers in ten PC9 cells. There were 8, 33, 54, 116, and 19 cases where the chromosomal number was 0, 1, 2, 3, and 4, respectively. The most frequent chromosomal number was three. We observed no Y chromosomes, as PC9 cells were derived from a female. There were no cases where the chromosomal number was more than 5.

| # | Cell 1 | Cell 2 | Cell 3 | Cell 4 | Cell 5 | Cell 6 | Cell 7 | Cell 8 | Cell 9 | Cell 10 |
|---|---|---|---|---|---|---|---|---|---|---|
| 1 | 3 | 4 | 3 | 3 | 3 | 4 | 4 | 3 | 3 | 3 |
| 2 | 3 | 3 | 3 | 3 | 3 | 3 | 3 | 3 | 2 | 3 |
| 3 | 3 | 3 | 2 | 3 | 3 | 3 | 3 | 3 | 3 | 3 |
| 4 | 1 | 1 | 1 | 1 | 1 | 1 | 1 | 1 | 1 | 1 |
| 5 | 3 | 3 | 2 | 3 | 3 | 3 | 3 | 3 | 3 | 3 |
| 6 | 3 | 2 | 3 | 3 | 3 | 3 | 3 | 3 | 3 | 3 |
| 7 | 3 | 3 | 3 | 3 | 3 | 2 | 3 | 3 | 3 | 3 |
| 8 | 2 | 2 | 2 | 2 | 1 | 2 | 2 | 3 | 2 | 2 |
| 9 | 3 | 3 | 4 | 3 | 4 | 3 | 3 | 3 | 4 | 4 |
| 10 | 3 | 3 | 3 | 3 | 3 | 3 | 3 | 3 | 3 | 3 |
| 11 | 4 | 4 | 4 | 4 | 4 | 4 | 4 | 4 | 4 | 4 |
| 12 | 3 | 3 | 3 | 3 | 3 | 3 | 3 | 3 | 3 | 3 |
| 13 | 2 | 2 | 2 | 1 | 2 | 2 | 2 | 2 | 2 | 2 |
| 14 | 1 | 1 | 2 | 1 | 1 | 1 | 2 | 1 | 1 | 2 |
| 15 | 3 | 3 | 3 | 3 | 3 | 3 | 3 | 3 | 3 | 3 |
| 16 | 3 | 1 | 3 | 3 | 3 | 3 | 2 | 3 | 3 | 3 |
| 17 | 3 | 4 | 0 | 3 | 3 | 3 | 3 | 3 | 3 | 3 |
| 18 | 2 | 3 | 2 | 2 | 2 | 1 | 2 | 3 | 2 | 2 |
| 19 | 2 | 2 | 2 | 2 | 2 | 2 | 2 | 2 | 3 | 2 |
| 20 | 3 | 3 | 4 | 3 | 3 | 3 | 3 | 2 | 3 | 3 |
| 21 | 0 | 0 | 1 | 0 | 0 | 0 | 0 | 1 | 0 | 1 |
| 22 | 1 | 1 | 2 | 1 | 1 | 1 | 2 | 1 | 1 | 1 |
| X | 2 | 2 | 2 | 2 | 2 | 1 | 2 | 2 | 2 | 2 |
| Y | 0 | 0 | 0 | 0 | 0 | 0 | 0 | 0 | 0 | 0 |

#### Supplementary Table 4. Oligonucleotide donor DNAs used in this study

| Name | Sequence |
|---|---|
| RBM20 R636S | ACAGATATGGCCCAGAAAGGCCGCGGTCTAGTAGTCCGGTGAGCCGGTCACTCTCCCCGA |
| ATP7B R778L | CATGCTCTTTGTGTTCATTGCCCTGGGCCTGTGGCTGGAACACTTGGCAAAGGTAACAGC |
| GRN R493X | CGGCTGGCTACACCTGCAACGTGAAGGCTTGATCCTGCGAGAAGGAAGTGGTCTCTGCCC |

#### Supplementary Table 5. gRNAs used in this study.

PAM Sequences are underlined. Bold letters indicate the sites of single nucleotide substitutions.

| Name | Sequence |
|---|---|
| RBM20 | GGTCT**C**GTAGTCCGGTGAGCCGG |
| ATP7B | GGGCC**G**GTGGCTGGAACACTTGG |
| GRN | GAAGGCT**C**GATCCTGCGAGAAGG |

#### Supplementary Table 6. First PCR primers for amplicon sequencing

The 3' side of the primer sequences, represented by lowercase letters, are gene-specific. The 5' side of the primers, represented by the uppercase letters, are the common adaptor sequences.

| Primer Name | Sequence (5' -> 3') |
|---|---|
| RBM20_FW-2 | TAACTTACGGAGTCGCTCTACGgtgtctgtgtgtgggtggggtggg |
| RBM20_RV-2 | GGATGGGATTCTTTAGGTCCTGagagctgcaggaggtgaagctgggag |
| ATP7B NGS FW-1 | TAACTTACGGAGTCGCTCTACGttattctctggtcatcctggtg |
| ATP7B NGS RV-2 | GGATGGGATTCTTTAGGTCCTGagcagctcttttctgaacctga |
| GRN-NGS FW1 | TAACTTACGGAGTCGCTCTACGataatgccattctgtgctcccttc |
| GRN-NGS RV1 | GGATGGGATTCTTTAGGTCCTGcactccacgtccttcacacc |

### Supplementary Table 7. Second PCR primer for amplicon sequencing

| Custom P5 index | Custom P7 index | Custom P5 index sequence | Custom P7 index sequence | Custom P5 adapter sequence (5'->3') |
|---|---|---|---|---|
| P01_PE1.0 | P01_PE2.0 | CAAGTGTTC | CAAGTGTTC | AATGATACGGCGACCACCGAGATCTACACTCTTTCCCTACACGACGCTCTTCCGATCTNNNNNCAAGTGTTCTAACTTACGGAGTCGCTCTACG |
| P02_PE1.0 | P02_PE2.0 | AGGACATTC | AGGACATTC | AATGATACGGCGACCACCGAGATCTACACTCTTTCCCTACACGACGCTCTTCCGATCTNNNNNAGGACATTCTAACTTACGGAGTCGCTCTACG |
| P03_PE1.0 | P03_PE2.0 | CACTAATGG | CACTAATGG | AATGATACGGCGACCACCGAGATCTACACTCTTTCCCTACACGACGCTCTTCCGATCTNNNNNCACTAATGGTAACTTACGGAGTCGCTCTACG |
| P04_PE1.0 | P04_PE2.0 | AGCCTGATG | AGCCTGATG | AATGATACGGCGACCACCGAGATCTACACTCTTTCCCTACACGACGCTCTTCCGATCTNNNNNAGCCTGATGTAACTTACGGAGTCGCTCTACG |
| P05_PE1.0 | P05_PE2.0 | TTACGCTAA | TTACGCTAA | AATGATACGGCGACCACCGAGATCTACACTCTTTCCCTACACGACGCTCTTCCGATCTNNNNNTTACGCTAATAACTTACGGAGTCGCTCTACG |
| P06_PE1.0 | P06_PE2.0 | ACTCTCCGT | ACTCTCCGT | AATGATACGGCGACCACCGAGATCTACACTCTTTCCCTACACGACGCTCTTCCGATCTNNNNNACTCTCCGTTAACTTACGGAGTCGCTCTACG |
| P07_PE1.0 | P07_PE2.0 | GTCGATGCA | GTCGATGCA | AATGATACGGCGACCACCGAGATCTACACTCTTTCCCTACACGACGCTCTTCCGATCTNNNNNGTCGATGCATAACTTACGGAGTCGCTCTACG |
| P08_PE1.0 | P08_PE2.0 | ACGGGAATT | ACGGGAATT | AATGATACGGCGACCACCGAGATCTACACTCTTTCCCTACACGACGCTCTTCCGATCTNNNNNACGGGAATTTAACTTACGGAGTCGCTCTACG |
| P09_PE1.0 | P09_PE2.0 | CGCGCCCAG | CGCGCCCAG | AATGATACGGCGACCACCGAGATCTACACTCTTTCCCTACACGACGCTCTTCCGATCTNNNNNCGCGCCCAGTAACTTACGGAGTCGCTCTACG |
| P10_PE1.0 | P10_PE2.0 | ACTAGTTTG | ACTAGTTTG | AATGATACGGCGACCACCGAGATCTACACTCTTTCCCTACACGACGCTCTTCCGATCTNNNNNACTAGTTTGTAACTTACGGAGTCGCTCTACG |
| P11_PE1.0 | P11_PE2.0 | AGTATTACA | AGTATTACA | AATGATACGGCGACCACCGAGATCTACACTCTTTCCCTACACGACGCTCTTCCGATCTNNNNNAGTATTACATAACTTACGGAGTCGCTCTACG |
| P12_PE1.0 | P12_PE2.0 | AGGTTGGGT | AGGTTGGGT | AATGATACGGCGACCACCGAGATCTACACTCTTTCCCTACACGACGCTCTTCCGATCTNNNNNAGGTTGGGTTAACTTACGGAGTCGCTCTACG |
| P13_PE1.0 | P13_PE2.0 | GTGAACCGA | GTGAACCGA | AATGATACGGCGACCACCGAGATCTACACTCTTTCCCTACACGACGCTCTTCCGATCTNNNNNGTGAACCGATAACTTACGGAGTCGCTCTACG |
| P14_PE1.0 | P14_PE2.0 | GCACAAAAC | GCACAAAAC | AATGATACGGCGACCACCGAGATCTACACTCTTTCCCTACACGACGCTCTTCCGATCTNNNNNGCACAAAACTAACTTACGGAGTCGCTCTACG |
| P15_PE1.0 | P15_PE2.0 | CTGTCTTCG | CTGTCTTCG | AATGATACGGCGACCACCGAGATCTACACTCTTTCCCTACACGACGCTCTTCCGATCTNNNNNCTGTCTTCGTAACTTACGGAGTCGCTCTACG |
| P16_PE1.0 | P16_PE2.0 | GACGCGACT | GACGCGACT | AATGATACGGCGACCACCGAGATCTACACTCTTTCCCTACACGACGCTCTTCCGATCTNNNNNGACGCGACTTAACTTACGGAGTCGCTCTACG |
| P17_PE1.0 | P17_PE2.0 | CAGCCCATA | CAGCCCATA | AATGATACGGCGACCACCGAGATCTACACTCTTTCCCTACACGACGCTCTTCCGATCTNNNNNCAGCCCATATAACTTACGGAGTCGCTCTACG |
| P18_PE1.0 | P18_PE2.0 | AGATATCTG | AGATATCTG | AATGATACGGCGACCACCGAGATCTACACTCTTTCCCTACACGACGCTCTTCCGATCTNNNNNAGATATCTGTAACTTACGGAGTCGCTCTACG |
| P19_PE1.0 | P19_PE2.0 | AATACGCAC | AATACGCAC | AATGATACGGCGACCACCGAGATCTACACTCTTTCCCTACACGACGCTCTTCCGATCTNNNNNAATACGCACTAACTTACGGAGTCGCTCTACG |
| P20_PE1.0 | P20_PE2.0 | TATCGTGCC | TATCGTGCC | AATGATACGGCGACCACCGAGATCTACACTCTTTCCCTACACGACGCTCTTCCGATCTNNNNNTATCGTGCCTAACTTACGGAGTCGCTCTACG |
| P21_PE1.0 | P21_PE2.0 | TCCTGGTAT | TCCTGGTAT | AATGATACGGCGACCACCGAGATCTACACTCTTTCCCTACACGACGCTCTTCCGATCTNNNNNTCCTGGTATTAACTTACGGAGTCGCTCTACG |
| P22_PE1.0 | P22_PE2.0 | GGCAGAGGA | GGCAGAGGA | AATGATACGGCGACCACCGAGATCTACACTCTTTCCCTACACGACGCTCTTCCGATCTNNNNNGGCAGAGGATAACTTACGGAGTCGCTCTACG |
| P23_PE1.0 | P23_PE2.0 | ACAATGGAG | ACAATGGAG | AATGATACGGCGACCACCGAGATCTACACTCTTTCCCTACACGACGCTCTTCCGATCTNNNNNACAATGGAGTAACTTACGGAGTCGCTCTACG |
| P24_PE1.0 | P24_PE2.0 | TAGTGGCCA | TAGTGGCCA | AATGATACGGCGACCACCGAGATCTACACTCTTTCCCTACACGACGCTCTTCCGATCTNNNNNTAGTGGCCATAACTTACGGAGTCGCTCTACG |
| P25_PE1.0 | P25_PE2.0 | ATCCTATCT | ATCCTATCT | AATGATACGGCGACCACCGAGATCTACACTCTTTCCCTACACGACGCTCTTCCGATCTNNNNNATCCTATCTTAACTTACGGAGTCGCTCTACG |
| P26_PE1.0 | P26_PE2.0 | CTGCTTAGC | CTGCTTAGC | AATGATACGGCGACCACCGAGATCTACACTCTTTCCCTACACGACGCTCTTCCGATCTNNNNNCTGCTTAGCTAACTTACGGAGTCGCTCTACG |
| P27_PE1.0 | P27_PE2.0 | CGTGGCTGC | CGTGGCTGC | AATGATACGGCGACCACCGAGATCTACACTCTTTCCCTACACGACGCTCTTCCGATCTNNNNNCGTGGCTGCTAACTTACGGAGTCGCTCTACG |
| P28_PE1.0 | P28_PE2.0 | GTAACTAAT | GTAACTAAT | AATGATACGGCGACCACCGAGATCTACACTCTTTCCCTACACGACGCTCTTCCGATCTNNNNNGTAACTAATTAACTTACGGAGTCGCTCTACG |
| P29_PE1.0 | P29_PE2.0 | GCTTACGAA | GCTTACGAA | AATGATACGGCGACCACCGAGATCTACACTCTTTCCCTACACGACGCTCTTCCGATCTNNNNNGCTTACGAATAACTTACGGAGTCGCTCTACG |
| P30_PE1.0 | P30_PE2.0 | GATGCAGTG | GATGCAGTG | AATGATACGGCGACCACCGAGATCTACACTCTTTCCCTACACGACGCTCTTCCGATCTNNNNNGATGCAGTGTAACTTACGGAGTCGCTCTACG |
| P31_PE1.0 | P31_PE2.0 | GGGCTACAA | GGGCTACAA | AATGATACGGCGACCACCGAGATCTACACTCTTTCCCTACACGACGCTCTTCCGATCTNNNNNGGGCTACAATAACTTACGGAGTCGCTCTACG |
| P32_PE1.0 | P32_PE2.0 | ATAGCACGC | ATAGCACGC | AATGATACGGCGACCACCGAGATCTACACTCTTTCCCTACACGACGCTCTTCCGATCTNNNNNATAGCACGCTAACTTACGGAGTCGCTCTACG |
| P33_PE1.0 | P33_PE2.0 | TATAACTCT | TATAACTCT | AATGATACGGCGACCACCGAGATCTACACTCTTTCCCTACACGACGCTCTTCCGATCTNNNNNTATAACTCTTAACTTACGGAGTCGCTCTACG |
| P34_PE1.0 | P34_PE2.0 | AAATGCGTT | AAATGCGTT | AATGATACGGCGACCACCGAGATCTACACTCTTTCCCTACACGACGCTCTTCCGATCTNNNNNAAATGCGTTTAACTTACGGAGTCGCTCTACG |
| P35_PE1.0 | P35_PE2.0 | CCATCCAGG | CCATCCAGG | AATGATACGGCGACCACCGAGATCTACACTCTTTCCCTACACGACGCTCTTCCGATCTNNNNNCCATCCAGGTAACTTACGGAGTCGCTCTACG |
| P36_PE1.0 | P36_PE2.0 | GTCTGCACC | GTCTGCACC | AATGATACGGCGACCACCGAGATCTACACTCTTTCCCTACACGACGCTCTTCCGATCTNNNNNGTCTGCACCTAACTTACGGAGTCGCTCTACG |

**Supplementary Table 8. Commands for the CRISPResso2 analysis used in this study.**

CRISPResso version (CRISPResso Batch mode): 2.0.45

**Commands used:**

/opt/conda/bin/CRISPResso -o /DATA/CRISPRessoBatch_on_batch --name **(File Name 1)** --needleman_wunsch_gap_extend -2 --aln_seed_count 5 --amplicon_name Reference --amplicon_seq **(Sequence 1)** --max_rows_alleles_around_cut_to_plot 50 --prime_editing_pegRNA_extension_quantification_window_size 5 --fastq_r1 **(File Name 2)** --quantification_window_size 1 --quantification_window_center -3 --trimmomatic_command trimmomatic --conversion_nuc_from C --min_bp_quality_or_N 30 --default_min_aln_score 30 --needleman_wunsch_gap_incentive 1 --min_paired_end_reads_overlap 10 --plot_window_size 20 --prime_editing_pegRNA_scaffold_min_match_length 1 --aln_seed_min 2 --aln_seed_len 10 --expected_hdr_amplicon_seq **(Sequence 2)** --needleman_wunsch_gap_open -20 --max_paired_end_reads_overlap 100 --guide_seq **(Sequence 3)** --dsODN **(Sequence 4)** --conversion_nuc_to T --flexiguide_homology 80 --flash_command flash --min_single_bp_quality 30 --exclude_bp_from_left 15 --needleman_wunsch_aln_matrix_loc EDNAFULL --min_average_read_quality 30 --min_frequency_alleles_around_cut_to_plot 0.2 --exclude_bp_from_right 15

File Name 1_Experiment Name

File Name 2_”File name”.fastq.gz

Sequence 1_Reference Sequence

Sequence 2_Excepted HDR Sequence

Sequence 3_gRNA Sequence

Sequence 4_Donor Sequence
